# A Novel *miR-4745*-KLC2 Axis Regulates Cancer Stem Cell Traits in Colorectal Cancer

**DOI:** 10.64898/2026.01.07.697660

**Authors:** Kuldeep Lahry, Pauline Barraja, Julie Ripoll, Océane Bouvet, Guillaume Pinna, Amandine Bastide, Stéphanie Mateus, Lisa Tellier, Kevin Bigot, Christy Matta, Elodie Pothin, Lucile Bansard, Cécile Dejou, Anne-Alicia Gonzalez, Eric Rivals, Julie Pannequin, Jean-Marc Pascussi, Alexandre David, Chris Planque

## Abstract

Advanced colorectal cancer (CRC) treatments often fail due to chemoresistance and tumor recurrence, primarily driven by cancer stem cells (CSCs), which play a significant role in therapy ineffectiveness. MicroRNAs (miRNAs) play a pivotal role in regulating numerous biological functions within CSCs. Through a gain-of-function miRNA screening, we identified miRNAs that regulate the ALDEFLUOR-positive CSC population in colon cancer, with *miR-4745* emerging as a promising inhibitor. Overexpression of *miR-4745* reduced CSC self-renewal, diminished chemoresistance, and prevented CSC enrichment after chemotherapy in patient-derived CRC models. Mechanistically, *miR-4745* directly targets the 3’UTR of kinesin-light chain 2 (*KLC2*) mRNA, a member of the kinesin superfamily associated with poor clinical outcomes across various cancers. siRNA-mediated *KLC2* downregulation reduced ALDH-positive CSCs and suppressed sphere formation, while *KLC2* overexpression promoted CSC traits. Additionally, we observed an inverse correlation between *KLC2* and *miR-4745* mRNA levels in different colon cancer cell culture settings. Notably, *KLC2* mRNA levels increased with local CRC progression and were correlated with poor overall survival in a cohort of over 1 000 stage I to IV colorectal cancer patients. Our study uncovers a novel *miR-4745*-KLC2 axis that regulates CSC properties and chemoresistance in CRC, offering a promising strategy to prevent relapse and improve clinical outcomes for patients.

## BACKGROUND

Colorectal cancer (CRC) is the most common cancer in men and the second most common in women worldwide (WHO). In the United States, colorectal cancer ranks as the third most common cancer diagnosis and the third leading cause of cancer-related mortality among both men and women (1). According to projections by the World Health Organization, the global impact of colorectal cancer is anticipated to grow substantially by 2040, with an estimated 63% rise in new cases and a 73% increase in yearly deaths. For localized CRC, surgery remains the standard of care, while advanced CRC is managed through a multimodal approach combining chemotherapy, targeted therapies, and immunotherapy. The primary chemotherapeutic agents include 5-fluorouracil (5-FU), irinotecan, and oxaliplatin, often administered in various combinations. However, these treatments fail in 90% of metastatic cases, primarily due to drug resistance and tumor recurrence driven by cancer stem cells (CSCs), a highly resilient subset of tumor cells (2). CSCs are defined by their expression of stem cell factors that confer stemness properties, including the ability to self-renew. Moreover, CSCs can initiate and sustain tumor growth in serial transplantation assays (3) . These cells exhibit increased resistance to treatments compared to other tumor cells (4,5) and often become more prevalent in CRC tumors following chemotherapy (6,7).

Considering the significant impact of CSCs on therapeutic failure in CRC patients, it is essential to develop and implement treatment approaches that increase the responsiveness of these cells to current therapies, thereby improving patient outcomes. However, the precise mechanisms driving the heightened therapeutic resistance of CSCs remain poorly understood. This resistance is multifactorial, arising from factors such as increased resistance to DNA damage, reduced apoptosis, enhanced autophagy, slow proliferation, and the activation of the WNT/β-catenin, Hedgehog, and Notch signaling pathways. Additionally, CSCs often express multidrug transporters such as ATP-binding cassette G2 (ABCG2) (8) and drug-metabolizing enzymes like aldehyde dehydrogenase 1A1 (ALDH1A1) (9). To identify CSCs in solid tumors, including CRC, flow cytometry techniques have been employed, targeting ABCG2 expression (Side Population, SP) and ALDH activity (ALDEFLUOR-positive cells) (10). Both SP cells and ALDH-positive cells exhibit increased resistance to cytotoxic drugs *in vitro* and *in vivo,* and their populations are enriched following chemotherapy (6).

Micro-RNAs (miRNAs) are small non-coding RNA molecules, typically 20-25 nucleotides in length, that regulate gene expression by binding to the 3’ untranslated region (3’UTR) of messenger RNA (mRNA), resulting in either translation inhibition or mRNA degradation. In cancer, miRNA expression is often dysregulated, and increasing evidence indicates that miRNAs can function as oncogenes or tumor suppressors. They play a crucial role in regulating gene expression related to various cancer-related processes, including tumor initiation, progression (11,12), and drug resistance (13).

Interestingly, miRNAs are increasingly recognized as pivotal regulators in the complex control of CSC fate. Multiple studies have highlighted their role in regulating CSC self-renewal across various cancers, including prostate cancer (14), breast cancer (15), and colon cancer (16). In colorectal CSCs specifically, overexpression of *miR-451* has been shown to reduce tumorigenicity, self-renewal, and resistance to irinotecan by downregulating the ABCB1 drug transporter (17). Moreover, *miR-137* suppresses the tumorigenic potential of colorectal cancer stem cells by directly targeting Musashi-1 (18), whereas *miR-140* contributes to resistance against methotrexate and 5-fluorouracil through its regulation of histone deacetylase 4 (*HDAC4*) (19).

We recently reported that *miR-148a* overexpression suppresses Pregnane X Receptor (PXR) signaling, which is crucial for the expression of chemoresistance genes such as *ALDH1A1* and *ABCG2* in CSCs. Our study further demonstrated that niclosamide induces *miR-148a*, thereby inhibiting PXR and preventing CSC chemoresistance and tumor recurrence, both *in vitro* and *in vivo* (16).

Given the critical role of CSCs in CRC chemoresistance and the impact of miRNAs on the CSC phenotype, we conducted a gain-of-function screen to identify miRNAs that regulate the ALDH-positive CSC population in CRC cells. This screen identified *miR-4745* as a novel and potent inhibitor of ALDH activity. Ectopic overexpression of *miR-4745* diminished the self-renewal and chemoresistance properties of colon CSCs and prevented their enrichment following chemotherapy. Subsequent analysis revealed that kinesin light chain 2 (*KLC2*) mRNA is a direct target of *miR-4745*. This microRNA binds to the 3′ untranslated region (3′UTR) of *KLC2* mRNA, resulting in decreased expression of the target transcript. Additionally, we observed an increase in *KLC2* expression with advancing cancer stages in colon cancer tissue samples.

These findings position *miR-4745* as a promising therapeutic candidate for reducing recurrence and drug resistance in CRC, while also uncovering a novel role for KLC2 in regulating colon CSCs.

## METHODS

### Cell lines and patient-derived tumor cell culture

CRC cell lines ((SW480 (RRID:CVCL_0546), SW620 (RRID:CVCL_0547), HT29 (RRID:CVCL_A8EZ), LS174T (RRID:CVCL_1384), HCT116 (RRID:CVCL_0291), LoVo

(RRID:CVCL_0399), and Colo205 (RRID:CVCL_0218)) were obtained from ATCC and cultured either in DMEM or RPMI medium (Gibco) supplemented with 10% FBS, or were cultured as spheroids in M11-defined medium using either ultra-low attachment flasks from Corning or flasks coated with Poly 2-hydroxyethyl methacrylate. M11 medium consisted of

DMEM/F12 (1:1) (Gibco), supplemented with N2, 3 mM Glutamine, 0.6% Glucose, 4µg/ml insulin (Sigma-Aldrich), 100U/ml Penicillin G, 100ug/ml Streptomycin, 10ng/ml hBasic-FGF (R&D Systems), and 20ng/ml hEGF (R&D Systems). Patient-derived colon cancer cells (CPP1, CPP6, CPP19, CPP25, CPP36, and CPP44) were established from CRC biopsies, while two circulating tumor cell lines (CTC44 and CTC45) were isolated from the blood of a patient with advanced metastatic colorectal cancer. These biological samples were provided by CHU-Carémeau (Nîmes, France) as part of a previously reported study (20,21), registered under ClinicalTrials.gov Identifier#NCT01577511 (First submitted: 2012-04-12; Last update posted: 2021-06-21). The study protocol was approved by the French Ethics Committee (Comité de Protection des Personnes, Sud Méditerrannée III). All relevant ethical regulations for research involving human participants were followed with written informed consent obtained from all patients. Cell cultures were maintained at 37°C in a humidified atmosphere with 5% CO_2_.

### Colon tumor tissue samples

RNA extracted from colon tumor tissue samples was obtained as part of the Clinical and Biological Database BCBCOLON (Institut du Cancer de Montpellier—Val d’Aurelle, France) registered under ClinicalTrials.gov Identifier #NCT03976960 (First submitted: 2019-06-05; Last update posted: 2025-02-12). Tissue samples were obtained from 41 patients at various stages of colon cancer: adenoma (n = 3), primary adenocarcinoma at stages I (n = 8), II (n = 11), III (n = 5), IV (n = 8) and metastases (n = 6). All patients were chemonaive.

The tumor samples were collected in accordance with French laws under the supervision of a designated investigator and registered with the French Ministry of Higher Education and Research (declaration number DC-2008–695). The research protocol received approval from the French Ethics Committee (CPP Sud Méditerranée III, reference number 2014.02.04) and was also authorized by the institutional translational research board (ICM-CORT-2018-28). All relevant ethical regulations for research involving human participants were followed, and written informed consent was obtained from all patients.

### Plasmid constructions

The dual sensor plasmid pmiR-sensor contains a bidirectional promoter that drives the expression of both AcGFP (a green fluorescent protein from *Aequorea coerulescens*) and the red fluorescent protein DsRed1 (22). In the 3’UTR of the GFP transcript, we inserted either a full-length 3’UTR of *KLC2* (at the *Xba*I restriction site) or two tandem copies of predicted *miR-4745* binding sites (between the *Apa*I and *Xba*I restriction sites). The sequences of the cloned regions are provided in **Supplementary Table S1**. GFP expression levels serve as a direct indicator of *miR-4745* activity, while DsRed1 is used for normalization. This dual expression system minimizes the effects of variable expression levels and nonspecific transcriptional regulation, allowing for ratiometric measurements (AcGFP/DsRed1). To create the pmiR-KLC2-3’UTR mutant #2, site directed mutagenesis was performed using the primers KLC2-3’UTR Mut2 Fp (CGGTACTACCCAAACCTCCCCTCGTCCCTCTTC) and KLC2-3’UTR Mut2 Rp (GACGAGGGGAGGTTTGGGTAGTACCGTGGACTG). The clones were confirmed via Sanger sequencing at Eurofins using the AcGFP Fp primer (CAACATCGAGGATGGCAG).

To create the KLC2-overexpressing plasmid (oe-KLC2), the *KLC2* coding sequence, lacking the 3’UTR *miR-4745* targeting sequence, was amplified from the pDONOR-KLC2 plasmid (Montpellier Genomic Collection MGC Facility) and inserted into the pCDNA3-3xFlag Cter plasmid at the *Hind*III and *Xba*I restriction sites.

### Transfection assay and oligonucleotides

The mirVana™ miRNA mimics, including *miR-4745-3p* (ref: 4464066, ID: MC21191), *miR-497-3p* (ref: 4464066, ID: MC12884), *miR-4723-5p* (ref: 4464066, ID: MC21507), *miR-4784* (ref: 4464066, ID: MC22683), and the mirVana microRNA Mimic Negative Control #1 (ref: 4464058), as well as the anti-miR^TM^ miRNA inhibitors, including anti-*miR4745-3p* (ref: AM17000, ID: AM21191) and the anti-miR™ miRNA Inhibitor Negative Control #1 (ref: AM17010) were purchased from Life Technologies. siRNA duplexes specific to *KLC2*, *HEY2*, *CCND3*, and *miR31HG* were purchased from Genecust, and their corresponding sequences are listed in **Supplementary Table S2**. Transfection of 50nM miRNA or siRNA duplexes was performed using Lipofectamine RNAiMax (Invitrogen) according to the manufacturer’s instructions. Additionally, transfection of 200 000 cells in a 6-well plate with 2µg of plasmid DNA was carried out using Lipofectamine 2000 (Invitrogen) for 72 hours, following the manufacturer’s protocol. For co-transfection experiments involving both a miRNA mimic and plasmid DNA, Lipofectamine 2000 was also used.

### miRNA extraction and quantification by real-time RT-PCR

miRNA was purified using the miRCURY^TM^ RNA isolation kit. First-strand cDNA synthesis was performed with the miRCURY LNA™ RT Kit (Qiagen), followed by real-time PCR amplification using the miRCURY LNA™ SYBR Green PCR Kit (Qiagen). Each sample was amplified in triplicate on an LC480 Real-Time PCR System (Roche). The expression levels of *miR-4745-3p* were calculated using the 2^−ΔΔCt^ method, with *miR-103a* used as the normalization control.

### RNA extraction and quantification by real-time PCR

Total RNA was extracted using the RNeasy Mini Kit (Qiagen) and treated with DNAse I to remove genomic DNA contamination. First-strand cDNA was synthesized using Superscript II (Invitrogen) and random hexamers, followed by real-time PCR for gene expression analysis. Each cDNA sample was analyzed in triplicate using SYBR Green chemistry (Roche) on a LightCycler 480 Real-Time PCR System (Roche). *GAPDH*, β*-actin*, *RPLO*, *18S ribosomal RNA*, and *RPL13* were used as endogenous controls. Gene expression fold changes were determined using the comparative Ct approach, specifically the 2^−ΔΔCt^ method. Primer sequences are provided in **Supplementary Table S3**.

### mRNA purification

mRNA was isolated from total RNA by performing two successive purifications with the GeneElute™ mRNA Purification Kit (Sigma), following the protocol provided by the manufacturer.

### Western blot

Cells were lysed using RIPA buffer supplemented with protease inhibitors (Roche) for total protein extraction. For western blotting, samples were separated on a 10% SDS-PAGE gel and transferred to a nitrocellulose membrane (Amersham). The antibodies utilized in this study included GAPDH (sc-32233, dilution 1:5,000; Santa Cruz), β-ACTIN (A-5441, dilution 1:10,000; Sigma-Aldrich), and KLC2 (17668-1-AP, dilution 1:1,000; Proteintech). Band intensities were quantified using Image Lab software (Bio-Rad).

### Proliferation assay

One thousand cells were seeded in triplicate in a 96-well plate and incubated for 24, 48, 72, and 96□hours. Cells were subsequently fixed in 10% trichloroacetic acid (TCA) at 4□°C for a minimum duration of 2 hours. After three washes with Milli-Q water, cells were stained with 0.4% Sulforhodamine B solution for 30□minutes at room temperature, followed by three additional washes with MilliQ water. Following resuspension in 10□mM Tris-Base, absorbance was recorded at a wavelength of 562□nm.

### Staining for flow cytometry and fluorescence-activated cell sorting (FACS)

CD44v6-APC (clone REA706, 130-111-238; Macs Miltenyi) antibody was incubated with 100 000 cells in PBS containing 5% FBS for 20 min at 4°C. The Aldefluor assay (Stem Cell Technologies) was performed according to the manufacturer’s instructions. ALDH-positive and ALDH-negative cells were identified by comparing the same sample treated with and without the ALDH inhibitor diethylaminobenzaldehyde (DEAB). The flow cytometry gating strategy for Aldefluor-stained samples involved initial staining using the ALEFLUOR assay, followed by SYTOX Blue Dead Cell Stain (Invitrogen). A stepwise gating strategy was employed to identify the primary cell population (SSC vs. FSC), isolate single cells (SSC-A vs. SSC-W), and select viable cells based on SYTOX Blue exclusion (SSC-A vs. SYTOX Blue fluorescence). Dead cells were excluded based on light scatter and SYTOX Blue Staining. DEAB-treated cells served as a negative control to set the gating threshold for identifying ALDH-positive subpopulation in test samples without DEAB. Cells were sorted using a FACSAria II instrument (BD), and the resulting data were processed and analyzed with Flowing Software version 2.5.1 (http://flowingsoftware.btk.fi/). ALDH activity was also assessed using the MACSQUANT analyzer (Miltenyi) with the same protocol.

For sphere formation assay experiments, live cells were sorted using FACSAria II (BD) based on negative labeling with SYTOX Blue or 7-AAD.

To experimentally validate the binding of *miR-4745* to the *KLC2* mRNA transcript, we employed a fluorescence-based dual reporter assay system using the pmiR-sensor plasmid. CPP1 cells were co-transfected with the pmiR-sensor plasmid containing the *KLC2* 3’UTR, and the *miR-4745-3p* mimic. GFP and DsRed1 expression levels were subsequently measured using a CytoFLEX LX flow cytometer (Beckman Coulter) at the MRI facility in Montpellier.

### In vitro cytotoxic treatment

Cells were plated at 200 000 cells per well in DMEM with 10% FBS in 6-well plates. After 24 hours, cells were exposed to FIRI (1X = 50μM 5-FU + 500nM SN38) or vehicle (0.1% DMSO), with 3 wells per condition. Following 72 hours of treatment, cells were washed twice with ice-cold PBS before RNA extraction.

### In vitro chemosensitivity assays

Cells were plated at 2 000 cells per well in DMEM with 10% FBS in 96-well plates. After 24 hours, they were treated with chemotherapeutic agents for 48 to 72 hours. Cell viability was evaluated using the CellTiter-Glo assay (Promega), and half-maximal effective concentration (EC_50_) values were determined using Prism9 software.

### Sphere formation assays

The percentage of cells forming spheres was determined by plating 100 or 200 live cells per well in M11 medium in ultra-low attachment 96-well plates. Live cells were sorted based on negative labeling with SYTOX Blue or 7-AAD. Spheres with a diameter exceeding 50 µm were counted.

### Polysome fractionation

This procedure was carried out as previously described (23). Briefly, 4 × 10^6^ CPP1 cells were seeded in Petri dishes (150 mm) and transfected 24 hours later with either the *miR-4745-3p* mimic or the mirVana microRNA Mimic Negative Control #1 (miCTRL). Following 48 hours of transfection, cells were exposed to 20 μg/ml emetine for 5 minutes at 37°C. Subsequently, they were rinsed twice with ice-cold PBS and harvested by scraping into ice-cold PBS. The cell suspension was centrifuged, resuspended in 1 mL of polysome lysis buffer, and homogenized by vigorous shaking with 1.4 mm ceramic spheres (Lysing matrix D MPBio) using a FastPrep machine (MPBio). The lysates were subsequently centrifuged at maximum speed for 10 minutes at 4°C.

The resulting supernatant (cytosolic fraction) was loaded onto a 15-50% sucrose gradient and centrifuged at 35 000 rpm for 2.5 hours at 4°C using a SW41 rotor (Beckman Coulter). Polysomes were separated and monitored in real-time using an optical density (OD) 254 nm UV spectrometer. Gradient fractionation was carried out using an ISCO density gradient fractionation system (Lincoln, NE), while continuously monitoring absorbance at 254 nm across the gradient depth.

### Transcriptome and Translatome analyses

Total RNA was isolated using TRIzol reagent (Invitrogen) following the manufacturer’s instructions. mRNA associated with 1 to 3 ribosomes (“Light polysomes”) and mRNA associated with more than 3 ribosomes (“Heavy polysomes”) were extracted using TRIzol LS (Invitrogen) according to the manufacturer’s instructions.

We prepared RNA-seq libraries using the NuGEN Universal Plus mRNA-seq kit (Tecan, Redwood City, CA, USA) according to the manufacturer’s instructions. Briefly, polyadenylated RNAs were selected using oligo-dT magnetic beads and chemically fragmented. First strand cDNAs were synthesized using random primers in the presence of actinomycin D to prevent spurious DNA-dependent synthesis. After second strand synthesis and end-repair, adapters including sequences used later in PCR amplification were ligated. The strand to be sequenced was purified and PCR amplified for 15 cycles. PCR products were purified using AMPure XP Beads (Beckman Coulter Genomics, Brea, CA, USA). The size distribution of the resulting libraries was monitored using a Fragment Analyzer (Agilent Technologies, Santa Clara, CA, USA) and the libraries were quantified using the KAPA Library quantification kit (Roche, Basel, Switzerland).

The libraries were denatured with NaOH, neutralized with Tris-HCl, and diluted to 300 pM. Clustering and sequencing were performed on a NovaSeq 6000 (Illumina, San Diego, CA, USA) using the single read 100nt protocol on 2 lanes of a flow cell S2.

Image analyses and base calling were performed in real time by the Real-Time Analysis software (Illumina). Demultiplexing was performed using Illumina’s conversion software (bcl2fastq 2.20).

### Bioinformatic analysis of transcriptome and translatome sequencing data

We describe the pipeline used to analyze RNA-seq data from the transcriptome (cytosolic fraction) and translatome (light and heavy polysomal fractions) under two conditions: control condition (CTL) and *miR-4745* overexpression (*miR-4745*). We refer to these three libraries as Cyto, Heavy and Light, and to the conditions as CTL and *miR-4745*. For each library, read quality was assessed using FastQC (v0.11.9, Babraham Institute, Cambridge, UK) and Illumina’s Sequencing Analysis Viewer (software SAV). FastqScreen (v0.15.1) was used to identify potential contamination. Reads were filtered using a quality threshold of Q35, and 12 bases at the 5’end were trimmed with Cutadapt v3.2. High-quality reads were then aligned to the Homo sapiens reference genome GRCh38 (primary assembly release 104) using STAR v2.7.9a (24), and annotated with the corresponding Ensembl annotation. Quantification of mapped reads was performed using featureCounts from subread v2.0.1 (25), and features with zero counts across all fractions were removed.

Statistical analyses were conducted using DESeq2 v1.30.1 (26) to identify differentially expressed genes (DEGs) between the two conditions for each fraction, with statistics computed using the Wald test. P-value adjustment for multiple testing was performed using the Benjamini-Hochberg method (27) to control the false discovery rate. Only genes with an adjusted p-value below 0.05 and a log fold change greater than one were selected for further analyses. Results from the cytosolic fraction were combined with those from the polysomal fraction to determine whether genes were regulated at the transcriptional level, translational level, or both. DEGs from pairwise comparisons (e.g., CTL-Heavy vs. *miR4745*-Heavy) were categorized using Pandas v1.1.5 into seven regulatory categories: no significant variation (background), regulation at the transcript level only (Transcription UP or Transcription DOWN), regulation at the mRNA translation level only (Translation UP or Translation DOWN), or regulation at both transcript and translation levels (total mRNA abundance UP or total mRNA abundance DOWN). Scatter plots illustrating gene variation and regulatory categories were generated using matplotlib v3.3.0 (28).

### miRNome-wide screening coupled with a miniaturized ALDEFLUOR assay

We conducted a miRNome-wide screening in the SW620 cell line using a miRNA overexpression (OE) library. The OE library contained miRNA mimics of endogenous miRNA duplexes (miRIDIAN® microRNA Mimics Library, Dharmacon/Horizon Discovery), designed to mimic 2 048 human miRNAs listed in miRBase v.19.0, as detailed previously (15). For screening, an automated reverse transfection method targeting pre-miRs was established using a robotic workstation fitted with a 96-well head probe (Nimbus, Hamilton). In short, pre-miRs were combined with Lipofectamine RNAiMAX (Life Technologies) and applied to collagen-coated, clear-bottom, black-walled 96-well culture plates (Costar Cat#3904). After 15 minutes of complexation, SW620 cells were seeded onto the lipoplexes (12 500 cells per well; final [pre-miR] = 25nM).

Three different controls were used in the OE screen: a commercial mixture of cytotoxic siRNAs as a positive transfection control (AllStars Maximal Death Control, Qiagen), a miRNA mimic against *miR-148a-3p* (phenotypic positive control, previously shown to downregulate the relative CSC proportion, Dharmacon/Horizon Discovery) and a mimic negative control (miRIDIAN® microRNA Mimic Negative Control, Dharmacon/Horizon Discovery). Three days post-transfection, the relative CSC proportion in SW620 cells treated with miRNA mimics was assessed using a previously described adaptation of the ALDEFLUOR assay (Stem Cell Technologies) for image acquisition and analysis in a 96-well format (29,30). Briefly, after removing the culture supernatants, 40 µL of an ALDEFLUOR/Hoechst mixture (5 µL of BAAA substrate per mL of ALDEFLUOR buffer + Hoechst 33342 [Sigma] at 50 µg/mL) were added to the culture wells. All plates included one control well that received N, N-diethylaminobenzaldehyde (DEAB), an inhibitor of ALDH. After 45 minutes of incubation at 37°C, the mixture was replaced with 50µL of ice-cold ALDEFLUOR buffer, and the plates were immediately imaged using a High Content Imaging device (Operetta HCS epifluorescence microscope, Perkin Elmer). During image acquisition, the plates were kept at 4°C. For each well, nine fields were captured at 10X magnification using two fluorescence channels: green for ALDEFLUOR (excitation: 470±10 nm; emission: 525±25 nm) and blue for Hoechst 33342 (ex: 380±20 nm; em: 445±35 nm). An automated algorithm was developed using Harmony 3.0 (Perkin Elmer) to quantify the relative CSC proportion. In summary, nuclear Regions of Interest (ROIs) were identified within the Hoechst channel, and the mean ALDEFLUOR signal, corrected for background, was quantified. Cells were defined as ALDH-positive when this signal exceeded the one quantified in the DEAB condition. The relative CSC proportion was computed as the number of ALDH-positive cells relative to the total number of cells.

### Data Preprocessing and Hit Selection

For each culture plate, the total cell amount and the percentage of ALDH-positive cells measured in sample wells were first normalized to the average values of the corresponding negative control (NEG) wells. The normalized results were labeled as *Relative Total Cell Amount* and *Relative % ALDH-positive cells*. A time□dependent decay of the ALDEFLUOR signal was observed, leading to a slight decrease in the *Relative % ALDH-positive cells* measured over the course of plate acquisitions. To account for and adjust this decay, a non-linear polynomial regression model was established for each plate to describe the relationship between the median *Relative % ALDH-positive cells* per column and the column index. For each column index, a correction factor was computed by dividing the plate’s median *Relative% ALDH-positive cells* by the predicted value from the model at the column index. These correction factors were subsequently applied to each column to normalize the individual *Relative % ALDH-positive cell* measurements. The corrected results were labeled as *Corrected Relative % ALDH-positive cells*. Analysis of the *Corrected Relative % ALDH-positive cells* revealed a non□Gaussian, long□tailed distribution in the sample population. To normalize this distribution, a Box–Cox transformation was applied to the population. The optimal λ coefficient for the Box–Cox transformation was determined by fitting a linear regression to quantile□to□quantile (QQ) plots, constructed from quantiles of the Box–Cox transformed *Corrected Relative % ALDH-positive cells* distribution plotted against quantiles of a corresponding theoretical Gaussian distribution. An optimal λ = 0.7 was determined to achieve the best linear fit. The normality of the Box–Cox transformed distribution was confirmed using a Kolmogorov–Smirnov test (p < 0.05), and the transformed results were labeled as *Box–Cox Corrected Relative % ALDH-positive cells.* The *Box–Cox Corrected Relative % ALDH-positive cells* sample results were then statistically scored using a Z□Score method. miRNA mimics were selected as hits if their absolute Z□Score was greater than 3.

### Statistical analysis

For each experiment, data are shown as mean ± S.E.M of n independent experiments (n = number of independent experiments). GraphPad Prism9 software was used for data analysis. The Mann Whitney test was used to analyse the difference between two groups of quantitative variables with alpha value set at 5%. *p<0.05; **p<0.005; ***p<0.001.

## RESULTS

### A miRnome-wide high content screen identifies *miR-4745* as a negative regulator of the ALDH-positive CRC population

We, and others, previously reported that the brightest subpopulation of CRC cells, identified using ALDEFLUOR^TM^ assay, a reporter for cellular ALDH enzymatic activity, is highly enriched in CSCs (10,21,29). These cells exhibit enhanced spheroid-forming efficiency and greater resistance to FIRI treatment. To investigate miRNAs involved in regulating colon CSCs, we conducted a miRNome-wide gain-of-function screen on a miniaturized ALDEFLUOR-based detection assay. This involved the systemic testing of a miRNA mimics library in the SW620 cell line (**Figure 1A**). SW620 was selected as the primary screening platform because it combines high technical suitability for automated high-content imaging with a biologically aggressive, metastatic phenotype characterized by a robust and stable ALDH+ cancer stem cell subpopulation. After 72 hours of miRNA transfection, CSCs were detected and quantified by measuring ALDEFLUOR-positive cells through high-content epifluorescence microscopy. The process involved identifying and counting cell nuclei stained with Hoechst dye and measuring cellular ALDEFLUOR intensities. ALDEFLUOR-positive cells were defined as those with intensities above the average intensity of cells treated with the ALDH inhibitor DEAB. The PARi platform optimized and miniaturized the ALDEFLUOR assay for 96-well plates, and developed automated procedures for image acquisition and analysis, as previously described for breast cancer cells (15). The screen revealed multiple miRNAs that significantly decreased ALDEFLUOR activity in SW620 cells (**Figure 1B; Supplementary Table S4**).

**Figure 1:**
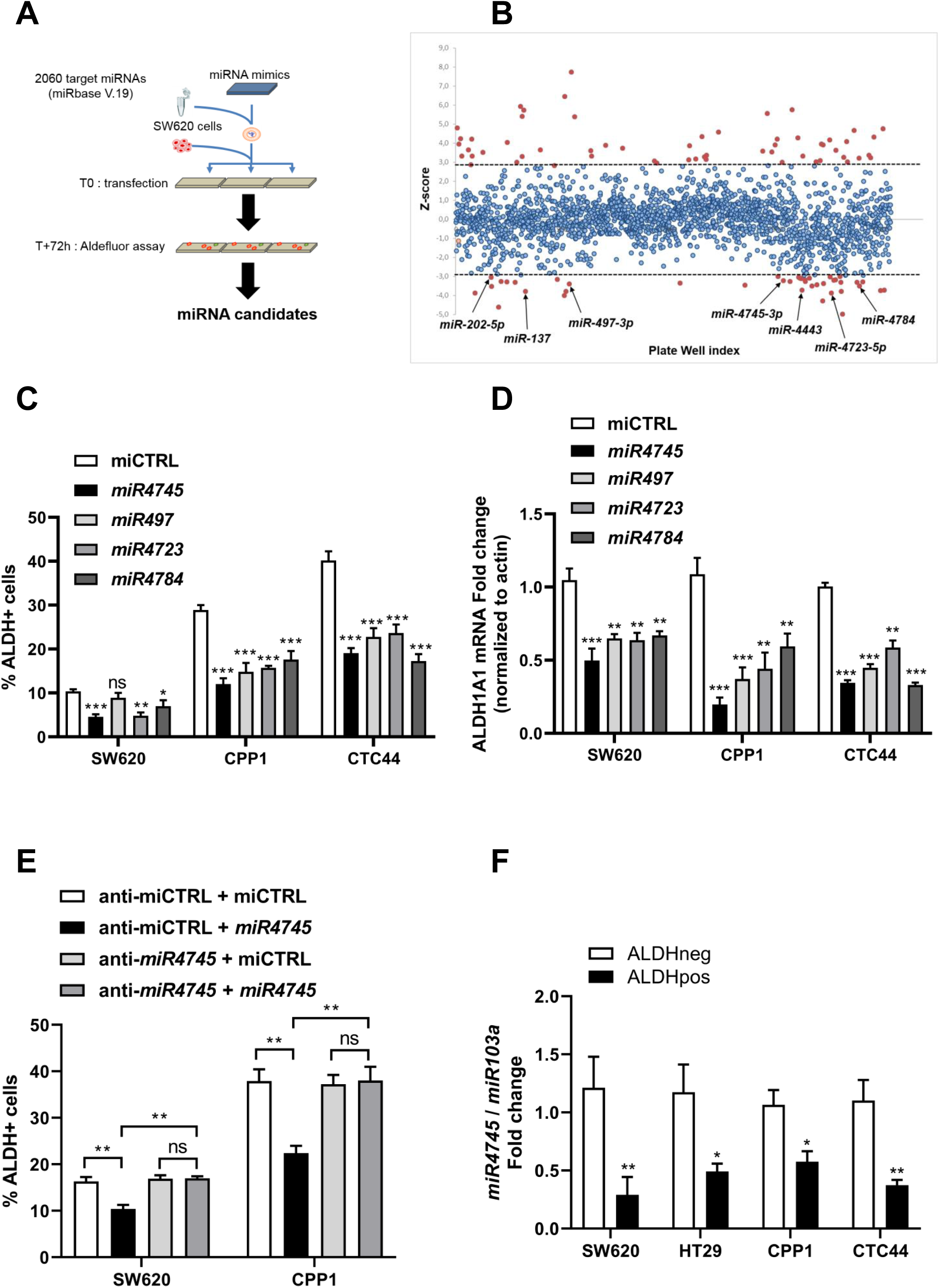
miRnome-wide High Content Screen identifies *miR-4745* as a negative regulator of the ALDH-positive CRC population. **(a)** Experimental design for the miRnome-wide gain-of-function screen employing pre-miRs as miRNA mimics in a scaled-down ALDEFLUOR detection assay. Following the individual transfection of miRNA mimics from a human miRNome-wide library into SW620 colon cancer cells, high-content screening was employed to measure the proportion of CSC (ALDEFLUOR-positive cells) in cells transfected with miRNA mimics or miRVana microRNA Mimic Negative Control #1 (miCTRL). **(b)** MA-plot of the high-content screening data, showing the Z-score of Aldefluor activity (mean of three replicates) in SW620 cells transfected with miRNA mimics from the human miRNome-wide library. The Z-scores are plotted against the plate well index, indicating the specific position of each miRNA mimic on the plate. **(c)** Percentage of Aldefluor-positive cells (% ALDH^+^ cells) in patient-derived CPP1 and SW620 colon cancer cells or circulating tumor cell line (CTC44) transfected for 72 hours with *miR-4745*, *miR-497*, *miR-4723*, *miR-4784* or miCTRL mimics. Data are expressed as mean ± SEM (n = 6). **(d)** RT-qPCR analysis of *ALDH1A1* mRNA expression in patient-derived CPP1 and SW620 colon cancer cells or circulating tumor cell line (CTC44) transfected for 72 hours with *miR-4745*, *miR-497*, *miR-4723*, *miR-4784* or miCTRL mimics. Data are expressed as mean ± SEM (n=3). Fold change is calculated relative to cells transfected with negative control mimic. **(e)** Percentage of Aldefluor-positive cells (% ALDH^+^ cells) in CPP1 or SW620 cells co-transfected for 72 hours with either anti-miR-4745 or anti-miCTRL, along with *miR-4745* or miCTRL mimics. Data are expressed as mean ± SEM (n = 4). **(f)** Expression levels of *miR-4745-3p* in patient-derived CPP1, SW620, and HT29 colon cancer cells, as well as in the circulating tumor cell line (CTC44), following cell sorting based on Aldefluor activity (ALDH-negative and ALDH-positive populations). Data are presented as mean ± SEM (n = 3) and are expressed as fold change relative to the ALDH-negative population. *, p < 0.05, **, p < 0.005, ***, p < 0.001, ns = not significant, Mann Whitney test.

Among the top miRNAs identified, miR-*137* (31), *miR-202-5p* (32) and *miR-4443* (33), have been previously reported to regulate multiple pathways involved in CSC maintenance and function, supporting the validity of our screening results. Subsequently, miRNAs were selected based on their relevance to CRC, CSC biology, and key phenotypic properties such as chemoresistance, self-renewal, and tumor initiation, with a particular focus on miRNAs that have been less extensively characterized but show promising potential.

We validated the activity of the selected miRNAs across three independent and complementary cellular models: the established SW620 cell line, patient-derived CRC cells from a primary tumor (CPP1), and patient-derived circulating tumor cell (CTC) lines (CTC44). CPP1 and CTC44 were specifically selected because they retain native genetic alterations and phenotypic heterogeneity that are often lost in long-established, immortalized cell lines. The inclusion of CTC44 is particularly significant as it provides a clinically relevant model of metastatic disease that retains the stem-like and chemoresistant features reflective of the original patient tumor. ALDEFLUOR activity was quantified by flow cytometry. Consistent with the screening findings, overexpression of *miR-4745*, *miR-497*, *miR-4723*, and *miR-4784* led to a significant decrease in the percentage of ALDH-positive cells relative to cells transfected with the negative control miRNA mimic (miCTRL) in all three cell lines (**Figure 1C**). This reduction in ALDH-positive cells correlated with decreased ALDH1A1 mRNA expression (**Figure 1D**). Among these candidates, *miR-4745* emerged as the most potent negative regulator of both the ALDH-positive cell population and ALDH1A1 mRNA expression, prompting further investigation into its effects.

To confirm that the reduction in ALDEFLUOR activity in CRC cells was specifically mediated by *miR-4745*, we used antagomiR-4745, a chemically modified RNA molecule designed to inhibit endogenous *miR-4745*. CPP1 and SW620 cells were co-transfected with either antagomiR-4745 (anti-*miR-4745*) to block miR-4745 activity, or a negative control (anti-miCTRL), along with *miR-4745* or miCTRL mimics. As shown in **Figure 1E**, the *miR-4745* mimic significantly decreased ALDEFLUOR activity in cells co-transfected with anti-miCTRL, but this effect was abolished in cells co-transfected with antagomiR-4745. This confirms that the reduction in ALDEFLUOR activity was specifically due to *miR-4745*, as blocking its activity with antagomiR negated the effect.

Finally, we assessed *miR-4745* expression in CSCs enriched from CRC cells using ALDEFLUOR cell sorting. We found that *miR-4745* expression was significantly lower in CSC populations (ALDH-positive cells) compared to non-CSC populations (ALDH-negative cells) sorted from SW620, CPP1, HT29, and CTC44 cell lines respectively (**Figure 1F**). These findings confirm that *miR-4745* expression or activity is downregulated in CSCs.

### Overexpression of *miR-4745* represses the CSC phenotype *in vitro*

To assess the functional effects of *miR-4745* modulation on CSC fate, CRC cell models, including patient-derived CTC lines representing metastatic and therapy resistant disease states, were transfected with a *miR-4745-3p* mimic. Initially, we confirmed that *miR-4745* overexpression does not affect cell growth, as cells transfected with the *miR-4745* mimic exhibited growth rates similar to those transfected with the control miRNA (miCTRL) (**Supplementary Figure 1A**). This indicates that any observed effects of *miR-4745* on the CSC phenotype are not attributable to changes in cell proliferation.

Subsequently, we assessed the presence of CD44v6-positive cells, a membrane marker linked to cancer stem cells and colorectal cancer metastasis (34), in CPP1 and CTC44 cells following *miR-4745* overexpression. We observed that *miR-4745* overexpression significantly reduced the percentage of CD44v6-positive cells (**Figure 2A**) and decreased CD44v6 mRNA expression (**Supplementary Figure 1B**) compared to cells transfected with the control miRNA (miCTRL).

**Figure 2:**
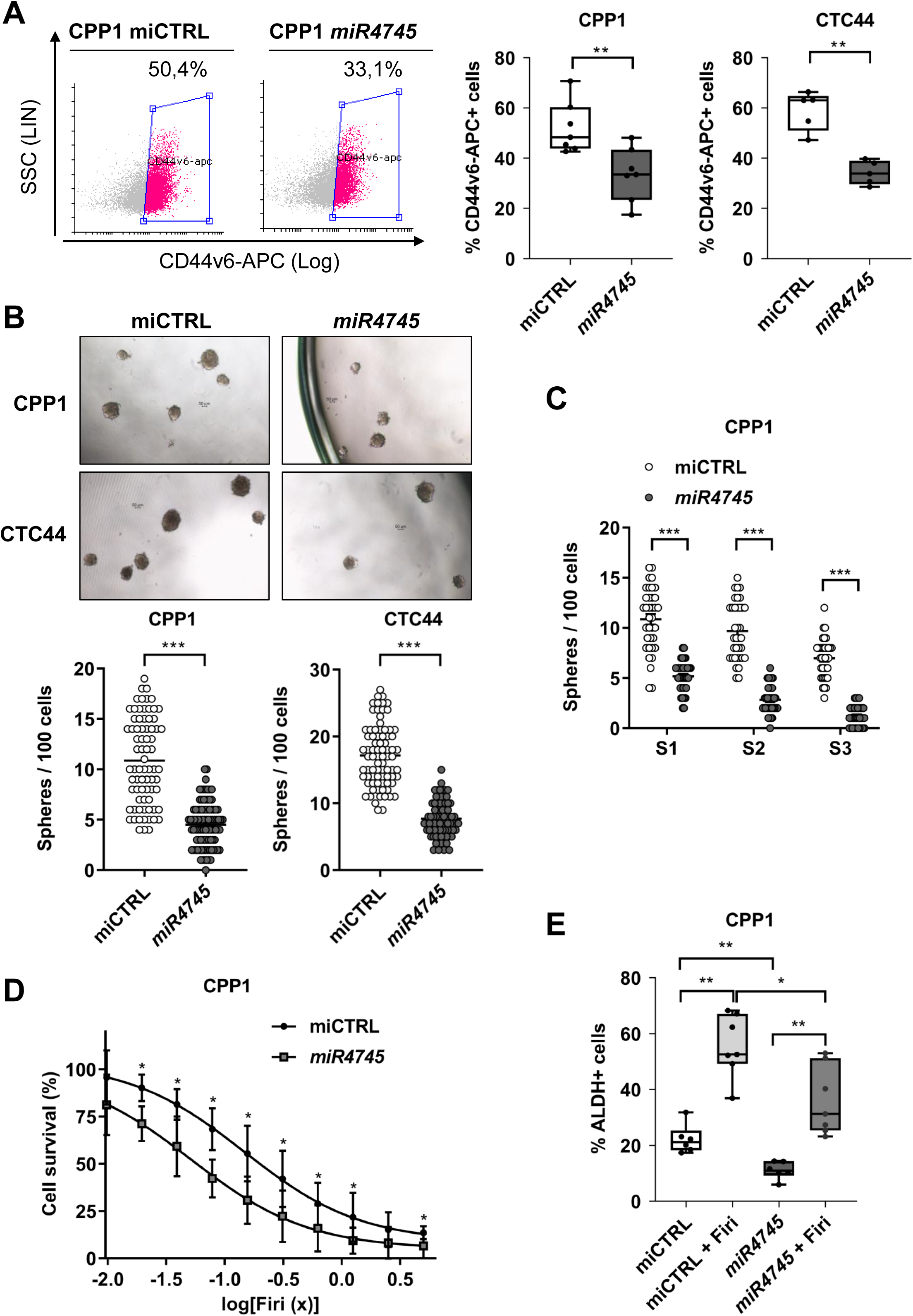
Overexpression of *miR-4745* represses the CSC phenotype *in vitro*. ***(a)*** Quantification of the CD44v6-APC-positive population in CPP1 (n=7) and CTC44 (n=5) cells transfected with *miR-4745-3p* (*miR4745*) or miRVana microRNA Mimic Negative Control #1 (miCTRL) mimics (right panel). Percentage of CD44v6-APC positive CPP1 cells is indicated in inset boxes for a representative experiment (left panel). **(b)** Representative images of tumorspheres (upper panel) and the percentage (lower panel) of sphere-forming cells in patient-derived CPP1 and circulating tumor (CTC44) colon cancer cells transfected with *miR-4745-3p* (*miR4745*) or miCTRL mimics. Data from individual replicates (n=15 per experiment) are presented, along with the mean ± SEM from six independent experiments. **(c)** Percentage of sphere-forming cells in patient-derived CPP1 colon cancer cells transfected with *miR4745* or negative control (miCTRL) mimics and, maintained as first-generation colonospheres (S1) or after several passages (S2, S3). Data from individual replicates (n=12 per experiment) are presented, along with the mean ± SEM from three independent experiments. **(d)** Percentage of surviving cancer stem cells (Aldefluor-positive) measured 48 hours following treatment with specified concentrations of FIRI (1X = 50 µM 5-FU + 500 nM SN38) *in vitro*. Sorted CPP1 Aldefluor-positive cells were first transfected with 50 nM *miR-4745-3p* (*miR-4745*) or miRVana microRNA Mimic Negative Control #1 (miCTRL) mimics, 24 hours prior to FIRI treatment. Data are expressed as mean ± SEM (n = 3). **(e)** Percentage of Aldefluor-positive cells (% ALDH^+^ cells) in CPP1 cells transfected with *miR-4745* or miCTRL mimics and then exposed for 72 hours to chemotherapy (Firi=5µM 5-FU + 50nM SN38). Data are expressed as mean ± SEM (n=6). *, p<0.05; **, p<0.005, ***, p < 0.001, Mann Whitney test.

We further evaluated the impact of *miR-4745* on the self-renewal capacity of CRC cells by assessing their ability to form tumorspheres. As shown in **Figure 2B** and **Supplementary Figure 1C**, cells transfected with the *miR-4745* mimic exhibited significantly lower spheroid-forming efficiency compared to those transfected with the negative control (miCTRL). To counteract the effects of *miR-4745* on sphere formation, we used antagomiR-4745. As depicted in **Supplementary Figure 1C**, the *miR-4745* mimic reduced the number of spheres in CPP1 and SW620 cells co-transfected with the antagomiR control sequence (anti-miCTRL + *miR4745*), but this effect was abolished in cells co-transfected with the *miR-4745* inhibitor (anti-*miR4745* + *miR4745*). Notably, the inhibitory effect of *miR-4745* extended beyond primary spheres to subsequent generations (**Figure 2C**), emphasizing its role in diminishing CSC self-renewal ability.

Next, we extended our investigation to chemoresistance, another hallmark of CSCs. We examined the effect of *miR-4745* overexpression on the survival of ALDH-positive patient-derived CRC cells treated *in vitro* with a standard CRC chemotherapy cocktail (FIRI = 5-fluorouracil [5-FU] + SN38, the active metabolite of irinotecan). As shown in **Figure 2D** and **Supplementary Figure 1D**, *miR-4745* transfection significantly decreased the survival of ALDH-positive cells after 48 hours of FIRI treatment, resulting in a four-fold decrease in EC_50_ (half maximal effective concentration) for CPP1 and CTC44 cells compared to those transfected with miCTRL (p < 0.05). Furthermore, **Figure 2E** and **Supplementary Figure 1E** illustrate that, although the proportion of ALDH-positive cells increased significantly following chemotherapy in cells transfected with miCTRL, this enrichment was notably reduced in cells transfected with *miR-4745*. These findings suggest that *miR-4745* sensitizes CSCs to chemotherapy. Taken together, these findings suggest that elevated *miR-4745* expression influences the cancer stem cell phenotype.

### *KLC2* is a potential target gene of *miR-4745*

Having established *miR-4745* as a regulator of CSC traits, our next objective was to identify its molecular targets. Given that miRNAs can modulate gene expression by degrading target mRNAs or inhibiting their translation, we conducted both transcriptome and translatome analyses in CPP1 cells overexpressing *miR-4745*. For transcriptome analysis, total RNAs were isolated from cytoplasmic lysates. For translatome analysis, mRNAs were isolated from polysome fractions of the same lysates, separated by sucrose gradient ultracentrifugation.

Overexpression of *miR-4745* did not alter the polysome profile in CPP1 cells, indicating that global protein synthesis remains unaffected (**Figure 3A**). To further investigate changes in translation, we sequenced mRNAs from both poorly translated fractions (light fractions: 1 to 3 ribosomes per mRNA) and highly translated fractions (heavy fractions: more than 3 ribosomes per mRNA) of the polysome profile. Overexpression of *miR-4745* in CPP1 cells resulted in selective modulation of the transcriptome, with 25 transcripts significantly upregulated and 20 downregulated (adjusted p-value < 0.05 and a fold change > 2 or < 0.5), highlighting a focused regulatory effect. Similarly, *miR-4745* overexpression induced targeted changes in mRNA translation, with 23 highly translated mRNAs upregulated and 17 downregulated, suggesting a refined role in translational control (**Figure 3B and Supplementary Tables S5 & S6**).

**Figure 3:**
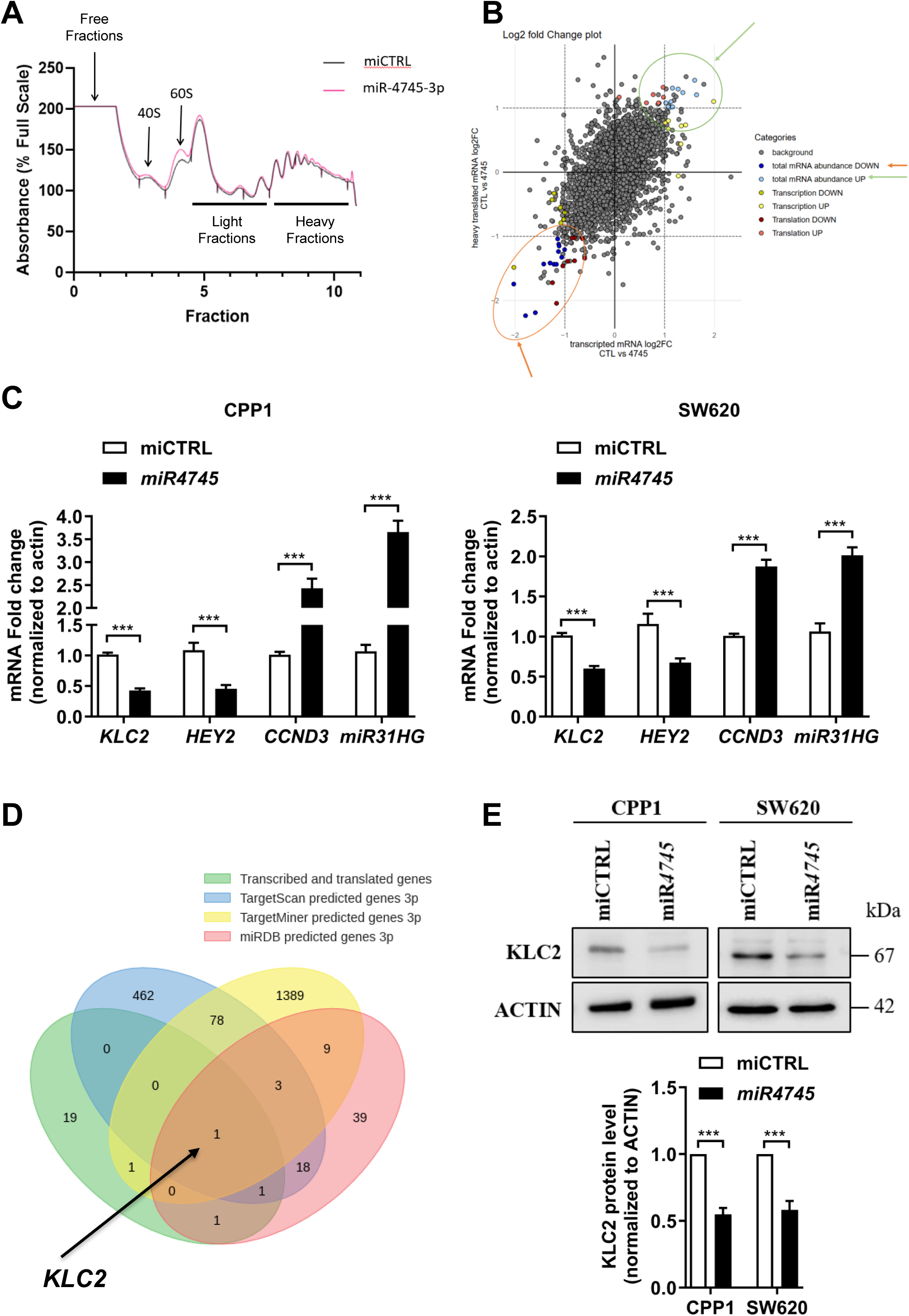
*KLC2* is a potential target gene of *miR-4745*. **(a)** Representative polysome profiling of CPP1 cells transfected with *miR-4745-3p* (*miR4745*) or miCTRL mimics for 72 hours. The profiles show peaks corresponding to the small (40S) and large (60S) ribosomal subunits, free fractions, light fractions (1 to 3 ribosomes per mRNA) and heavy fractions (more than 3 ribosomes per mRNA). This experiment was repeated three times. **(b)** Scatter plot showing the differential distribution of the log2 fold change of RNA from heavy fractions (y-axis) versus the log2 fold change of RNA from free fractions (x-axis) in CPP1 cells transfected with *miR-4745* or miCTRL mimics. **(c)** mRNA expression levels of potential *miR-4745* target genes were assessed in CPP1 and SW620 cells transfected with *miR-4745-3p* (*miR4745*) or miRVana microRNA Mimic Negative Control #1 (miCTRL) mimics for 72 hours. Data are presented as mean ± SEM (n = 3) and are expressed as fold change relative to cells transfected with the miCTRL mimic. (**d**) Venn diagram showing the intersection of the sets of *miR-4745* target genes predicted by TargetScan (blue), TargetMiner (yellow), miRDB (red), and the set of differentially expressed genes (DEGs) regulated at both transcriptional and translational levels (green - UP or DOWN). The intersection of all four sets contains a single gene, *KLC2*. (**e**) Representative Western blot analysis (upper panel) and quantification (lower panel) of KLC2 protein levels in patient-derived CPP1 and SW620 colon cancer cells transfected for 72 hours with either *miR-4745-3p* (*miR4745*) or miCTRL mimics. Data are presented as mean ± SEM (n=4). ***, p<0.001, Mann Whitney test.

We then validated the RNA sequencing results by analyzing the expression of two of the most downregulated genes (*KLC2* [Kinesin Light Chain 2] and *HEY2* [Hes Related Family BHLH Transcription Factor With YRPW Motif 2]) and two of the most upregulated genes (*CCND3* [Cyclin D3] and *MIR31HG* [miR-31 host gene]) at both transcriptional and translational levels, following transient overexpression of *miR-4745* in SW620 and CPP1 cell lines (**Figure 3C**).

Within each category of significantly differentially expressed genes (DEGs) (see section *Bioinformatic analysis of transcriptome and translatome sequencing data* in Materials and Methods), we searched for *miR-4745* target sites using TargetScan, TargetMiner, and the miRDB database. Since *miR-4745* target sites are preferentially located in the 3’ UTR of messenger RNAs, we computed the intersection of DEGs in each category with the set of predicted *miR-4745* target genes from each tool or database. As shown in the Venn diagram in **Figure 3D**, only one differentially expressed gene was predicted by all three approaches to have a *miR-4745* target site in its 3’ UTR. This gene was *KLC2*, which is regulated at both mRNA and protein levels. The downregulation of KLC2 at the protein level with *miR-4745* overexpression was confirmed by immunoblotting in CPP1 and SW620 cells (**Figure 3E**).

### *MicroRNA-4745* targets *KLC2* mRNA by binding to a specific sequence within its 3’untranslated region (3’UTR)

To experimentally confirm the binding of *miR-4745* to *KLC2* mRNA, we employed a fluorescence-based dual reporter assay system using a pmiR-sensor plasmid (16). This plasmid features a bidirectional promoter that drives the simultaneous expression of GFP from *Aequorea coerulescens* (AcGFP) and the red fluorescent protein DsRed1 (22). The GFP’s 3’UTR was modified to include the full-length 3’UTR of *KLC2* (*KLC2*-3’UTR), which contains a binding site for *miR-4745* (**Figure 4A**). Thus, the level of GFP reflects *miR-4745* activity, while DsRed1 expression normalizes for expression variability and nonspecific transcriptional regulation, providing a ratiometric measure (GFP/DsRed).

**Figure 4:**
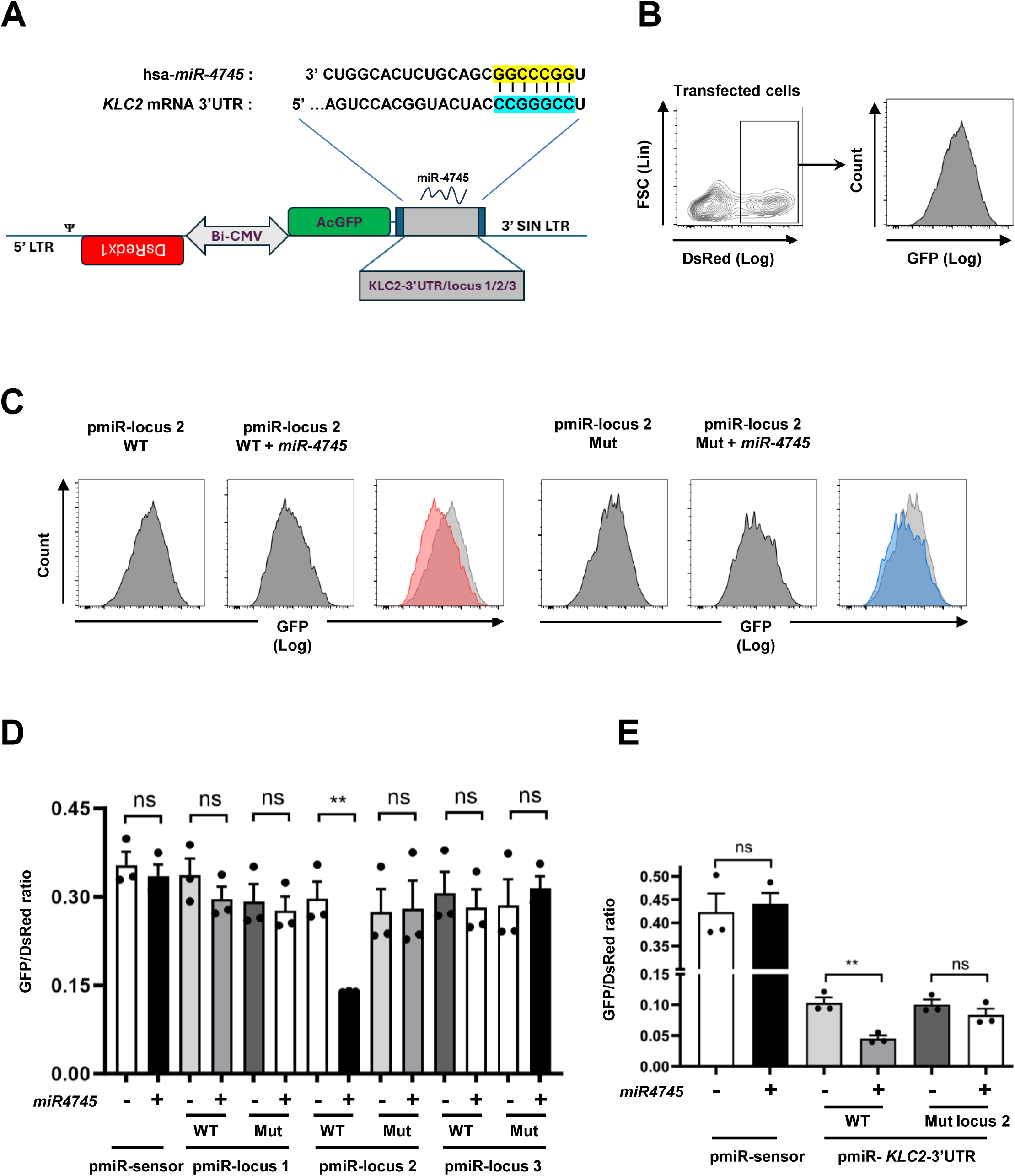
*MicroRNA-4745* targets *KLC2* mRNA by binding to a specific sequence in its 3’untranslated region (3’UTR). **(a)** The schematic illustration depicts the dual-reporter pmiR-sensor plasmid, in which the full-length *KLC2*-3’UTR or one of its three predicted binding sites (loci #1, #2, and #3) for *miR-4745*, is inserted into the 3’UTR of AcGFP (GFP). (**b**) Flow cytometry was used to measure the fluorescence intensities of GFP and DsRed in DsRed-positive cells. (**c**) Flow cytometry analysis (count versus GFP) was conducted on CPP1 cells co-transfected with a pmiR-sensor plasmid containing two copies of either the wild-type (WT) or mutated (Mutant) predicted binding site within the KLC2-3’UTR (pmiR-locus #2 WT or mutant), along with a *miR-4745-3p* mimic. (**d**) The GFP/DsRed ratio (geometric mean of intensities) was analyzed for CPP1 cells co-transfected with the pmiR-sensor plasmid containing two copies of the predicted binding sites within the KLC2-3’UTR or their mutant versions (sequences listed in **Supplementary Table S4**), along with a *miR-4745-3p* mimic. (**e**) The GFP/DsRed ratio (geometric mean of intensities) was analyzed for CPP1 cells co-transfected with the pmiR-sensor plasmid containing the full-length KLC2-3’UTR or its mutated version at locus #2, along with a *miR-4745-3p* mimic. Data are expressed as mean ± SEM (n=3). **, p<0.005, Student’s t-test.

CPP1 cells were co-transfected with the miR-sensor plasmid containing the *KLC2* 3’UTR (pmiR-*KLC2*-3’UTR), with or without a *miR-4745-3p* mimic. GFP and DsRed expression levels were then measured using flow cytometry (**Figures 4B** to **4E**). As shown in **Figure 4E**, co-transfection with the *miR-4745-3p* mimic specifically led to a decrease in GFP expression in cells transfected with the pmiR-*KLC2*-3’UTR compared to other conditions, indicating that *miR-4745-3p* binds to the *KLC2* 3’UTR.

To pinpoint the exact binding location of *miR-4745* within the *KLC2* 3’UTR, we used TargetScan, an online database that predicts miRNA binding sites in mammalian mRNAs transcripts. This analysis identified three 7-mer binding sites (seed sequence - CCGGGCC) within the *KLC2* 3’UTR that match the seed region (position 2-8 of the mature miRNA) of *miR-4745* (**Supplementary Figure 2**). The locations were identified as loci 1, 2, and 3. Two copies of each locus were cloned into the 3’UTR of GFP in the pmiR-sensor plasmid, and mutant versions of these loci (CGGGCC to CAAACC) were generated to disrupt *miR-4745* binding. As shown in **Figures 4C** and **4D**, GFP intensity significantly decreased when the wild-type locus 2 construct was co-transfected with *miR-4745*, but this effect was rescued when the binding site at locus 2 was mutated. No significant effects were observed at the other loci (**Figure 4D)**. This data confirms that *miR-4745* specifically binds to locus 2 within the *KLC2* 3’UTR and downregulates *KLC2* expression. Furthermore, mutating the same locus in the pmiR-KLC2-3’UTR plasmid also rescued the effect of the *miR-4745* mimic (**Figure 4E**). An overall decrease in fluorescence was seen when the full-length *KLC2* 3’UTR was cloned into the 3’UTR of GFP compared to the pmiR-sensor, which may be due to its larger size (∼1 kb). Overall, these findings validate the specific binding of *miR-4745* at locus 2 within the *KLC2* 3’UTR, leading to the downregulation of *KLC2* expression.

### Downregulation of *KLC2* reduces the colorectal CSC phenotype

Our research has established that *miR-4745* modulates *KLC2* expression, suggesting that the *miR-4745*-*KLC2* axis may be involved in the CSC phenotype. To elucidate the direct role of KLC2 in this process, CPP1 and SW620 cells were transfected with siRNA targeting *KLC2* (si*KLC2*) **(Supplementary Figures 3A** and **3B**), and ALDEFLUOR activity was assessed. Consistent with the effects observed with the *miR-4745* mimic, the knockdown of *KLC2* resulted in a significant reduction in the ALDH-positive cell population in both CPP1 and SW620 cells compared to controls transfected with the non-targeting siRNA (siCTRL) (**Figure 5A**).

**Figure 5:**
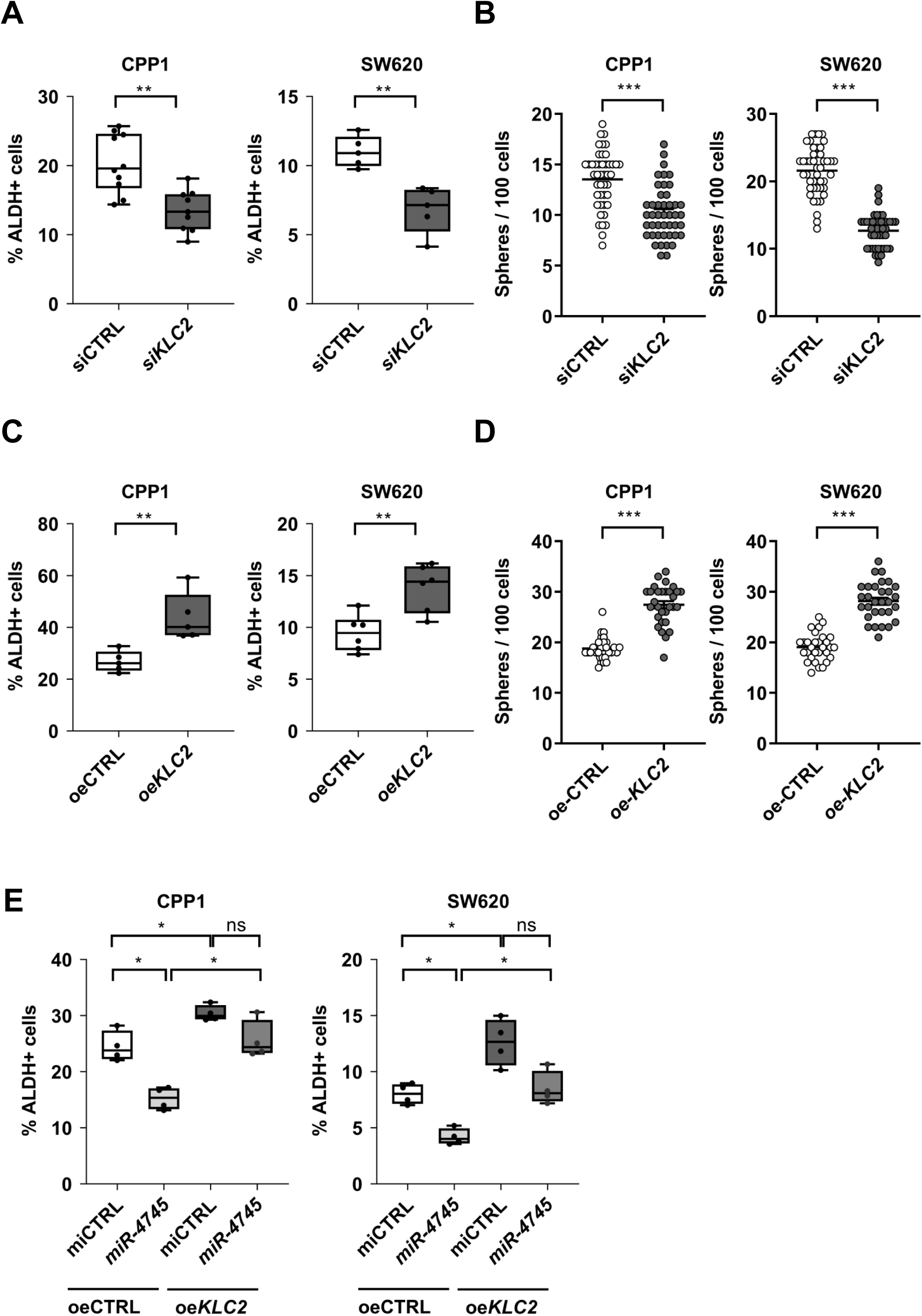
*KLC2* inactivation inhibits colorectal CSC phenotype. **(a)** Percentage of Aldefluor-positive cells (% ALDH^+^ cells) in CPP1 or SW620 cells transfected for 72 hours with 50nM of control (siCTRL) or KLC2-specific (*siKLC2*) siRNA. Data are expressed as mean ± SEM (n = 9 for CPP1 and n = 5 for SW620). **(b)** Percentage of sphere-forming cells among sorted live (Sytox Blue-negative) CPP1 and SW620 cells transfected for 72 hours with 50nM of control (siCTRL) or KLC2-specific (siKLC2) siRNA. Data are expressed as mean ± SEM (n=3 experiments with 15 individual replicates per experiment). **(c)** Percentage of Aldefluor-positive cells (% ALDH^+^ cells) in CPP1 or SW620 cells transfected for 72 hours with either a control plasmid (oe-CTRL) or a plasmid overexpressing *KLC2* (oeKLC2). Data are expressed as mean ± SEM, n = 5. **(d)** Percentage of sphere-forming cells among sorted live (Sytox Blue-negative) CPP1 and SW620 cells transfected for 72 hours with either a control plasmid (oe-CTRL) or a plasmid overexpressing KLC2 (oeKLC2). Data are presented as mean ± SEM, n = 3 (10 individual replicate values per experiment). **(e)** Percentage of Aldefluor-positive cells (% ALDH^+^ cells) in CPP1 or SW620 cells co-transfected for 72 hours with either a control plasmid (oe-CTRL) or a plasmid overexpressing *KLC2* lacking the 3’UTR *miR-4745*-targeting sequence (oeKLC2), along with miR-4745 or miCTRL mimics. Data are presented as mean ± SEM (n = 4). *, p < 0.05; **, p < 0.005; ***, p < 0.001; ns = not significant, Mann Whitney test.

To evaluate the proportion of CSCs within the tumor population, we performed a sphere formation assay. As illustrated in **Figure 5B**, *KLC2* knockdown reduced the ability of CPP1 and SW620 cells to form tumorspheres *in vitro* by approximately 40% to 50%, with no change in cell proliferation (**Supplementary Figure 3C**). Conversely, transient overexpression of *KLC2* in CRC cells (**Supplementary Figure 3D**) enhanced the stem cell phenotype, as evidenced by increased ALDEFLUOR activity and improved tumorsphere formation capacity *in vitro* (**Figures 5C** and **5D**).

To demonstrate that the effect of *miR-4745* on the CSC phenotype is mediated through *KLC2*, SW620 and CPP1 cells were co-transfected with either a *KLC2* overexpression plasmid construct lacking the 3’UTR *miR-4745* binding site (oeKLC2) or a control plasmid (oeCTRL), along with *miR-4745* or miCTRL mimics. While ALDEFLUOR activity decreased in cells co-transfected with the control plasmid and *miR4745* mimic, the inhibitory effect of *miR-4745* on ALDH-positive cells was counteracted in cells co-transfected with the cDNA construct lacking the 3’UTR *miR-4745* binding site (**Figure 5E**). This rescue experiment supports the hypothesis that *KLC2* downregulation is a crucial mechanism through which *miR-4745* exerts its effects.

### *miR-4745* negatively correlates with *KLC2* expression in CRC cells

Given our demonstration of the direct post-transcriptional regulation of *KLC2* by *miR-4745* in colon cancer cells, we expected to observe an inverse relationship between *miR-4745* and *KLC2* levels in colorectal cancer cells. RT-qPCR analysis revealed a significant negative correlation between *KLC2* mRNA and *miR-4745* (r = -0.716, p = 0.0027) across a panel of CRC cell lines (SW480, SW620, HT29, LS174T, HCT116, Lovo, and Colo205), as well as in patient-derived CRC cells from primary tumors (CPP1, CPP25, CPP44), metastatic tumors (CPP6, CPP19, CPP36), and circulating tumor cells (CTC44, CTC45) (**Figure 6A**).

**Figure 6:**
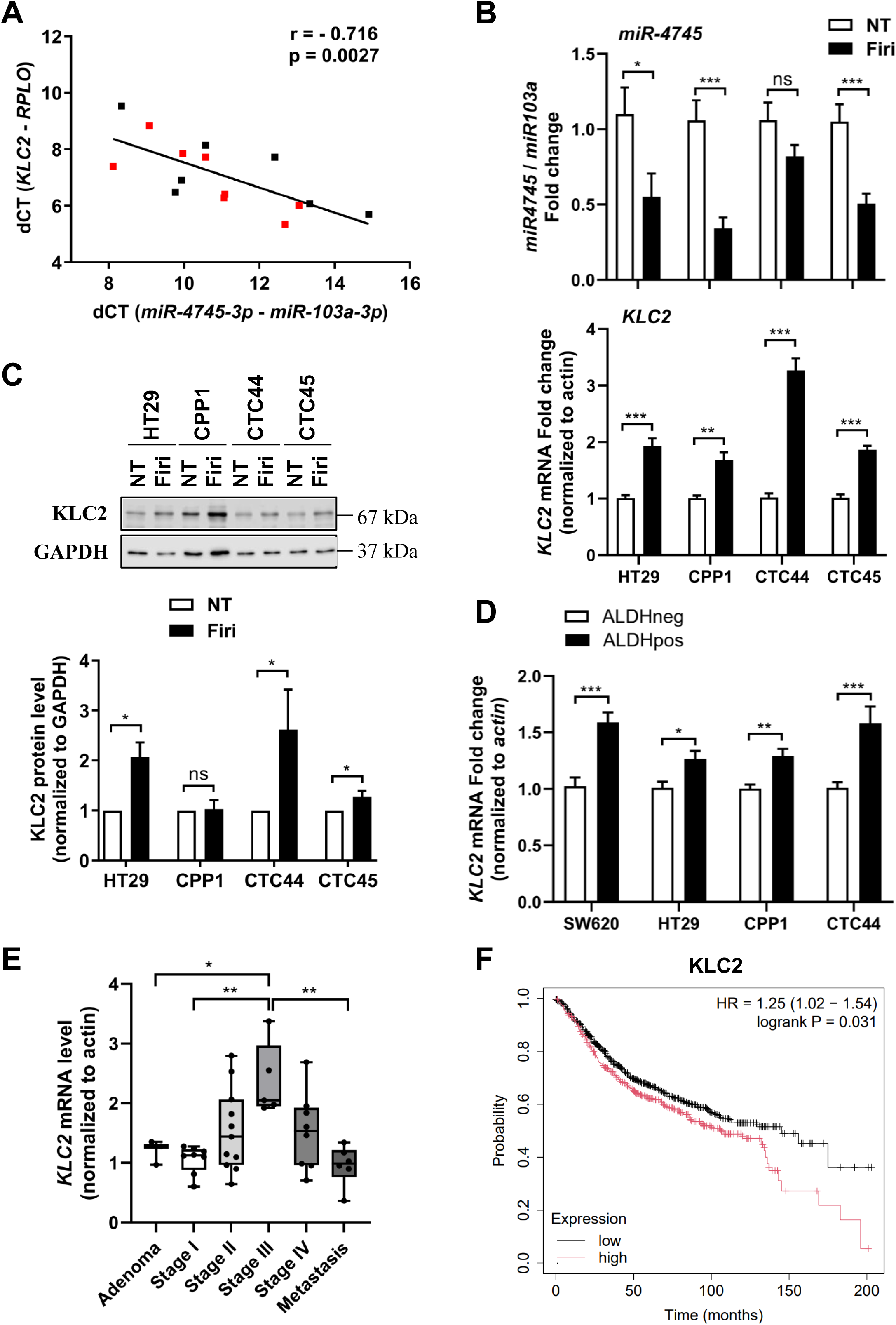
*KLC2* is inversely correlated with *miR-4745-3p* in CRC and is upregulated under chemotherapy treatment and with local CRC progression. **(a)** Inverse correlation between *KLC2* and *miR-4745-3p* expression in CRC cell lines obtained from ATCC (SW480, SW620, HT29, LS174T, HCT116, Lovo, and Colo205, black squares) and patient-derived CRC cells established in the IGF laboratory, including those from primary tumors (CPP1, CPP25, CPP44), metastatic tumors (CPP6, CPP19, CPP36), and blood (circulating tumor cells, CTC44, CTC45, red squares). Expression levels were measured by RT-qPCR. *KLC2* expression (mean of each technical replicate) was normalized to RPL0 (Ribosomal protein L0), while *miR-4745-3p* expression was normalized to *miR-103a-3p*. (**b**) *miR-4745-3p* (upper panel) and *KLC2* (lower panel) mRNA expression in HT29, CPP1, CTC44, and CTC45 colon cancer cells after 72 hours of chemotherapy treatment (Firi = 5 µM 5-FU + 50 nM SN38). Data are presented as mean ± SEM (n=3), showing fold change compared to untreated cells (NT). (**c**) Representative Western blot analysis (upper panel) and quantification (lower panel) of KLC2 protein expression levels in HT29, CPP1, CTC44, and CTC45 colon cancer cells after 72 hours of chemotherapy treatment (Firi = 5 µM 5-FU + 50 nM SN38). Data are presented as mean ± SEM (n=4), showing fold change compared to untreated cells (NT). **(d)** Expression levels of *KLC2* in patient-derived CPP1, SW620, and HT29 colon cancer cells, as well as in the circulating tumor cell line (CTC44), following cell sorting based on Aldefluor activity (ALDH-negative and ALDH-positive populations). Data are presented as mean ± SEM (n = 3) and are expressed as fold change relative to the ALDH-negative population. (e) *KLC2* mRNA expression in colon cancer tissue samples from 41 patients at various stages of cancer: adenoma (n = 3), primary adenocarcinoma at stages I (n = 8), II (n = 11), III (n = 5), IV (n = 8) and metastases (n = 6). Each dot represents an individual value. *KLC2* mRNA expression was normalized to actin. *, p < 0.05; **, p < 0.005; ***, p < 0.001, Mann Whitney test (**b** to **d**) or two-way ANOVA (Tukey’s multiple comparisons test, **e**). **(f)** Kaplan-Meier estimates of overall survival probability based on *KLC2* expression levels in a cohort of 1 061 stage I to IV colorectal cancer patients (https://kmplot.com/analysis/index.php?p=service&cancer=colon). Patients with low *KLC2* expression are depicted in black, while those with high *KLC2* expression are shown in red. The p-value from the likelihood ratio test is indicated.

Furthermore, *miR-4745* expression was significantly lower in CSCs enriched from CRC cells treated with chemotherapy (FIRI = 5-FU + SN38) compared to non-CSCs. This pattern was consistent in both ATCC-derived CRC cells (HT29) and patient-derived CRC cells (CPP1, CTC44, CTC45) under CSC-enriching conditions (**Figure 6B**). In contrast, *KLC2* mRNA and protein levels significantly increased following FIRI treatment, exhibiting an expression pattern opposite to that of *miR-4745* (**Figures 6B** and **6C**).

Additionally, *KLC2* transcript levels were elevated in ALDH-positive cells across all four CRC lines tested, suggesting that the downregulation of *miR-4745* contributes to the preferential expression of *KLC2* in these cells (**Figure 6D**).

### Differential expression of *miR-4745* and *KLC2* in human CRC samples

To further explore the expression patterns of *miR-4745* and *KLC2* in human colorectal cancer samples, we analyzed data from The Cancer Genome Atlas (TCGA) and dbDEMC databases. In the TCGA dataset, *miR-4745-3p* expression was found to be marginally lower in colon adenocarcinoma samples compared to non-cancerous tissues (n=103 vs. n=3, p = 0.06), although this difference was not statistically significant, possibly due to the small size of the non-cancerous group. Conversely, analysis of the dbDEMC dataset demonstrated a significant decrease in *miR-4745-3p* expression in colon adenocarcinoma samples (n=444) relative to non-cancerous controls (n=8, p = 0.00125). Additionally, we measured *KLC2* mRNA expression in colon cancer tissues from 41 patients across different stages of CRC: adenoma (n=3), stage I (n=8), stage II (n=11), stage III (n=5), stage IV (n=8), and metastasis (n=6). *KLC2* expression showed a significant increase with early local progression of CRC, suggesting its involvement in the early stages of tumor development (**Figure 6E**). However, a reduction in *KLC2* mRNA levels was observed in stage IV and metastatic samples, where cancer cells had spread from their primary site, suggesting that the gene may be subject to microenvironment-dependent regulation or reprogramming in advanced disease stages (35,36) (**Figure 6E**).

In addition, we sought to determine whether *KLC2* expression correlates with clinical outcomes in colorectal cancer patients. We performed a Kaplan-Meier analysis of *KLC2* mRNA expression in a cohort of stage I to IV CRC patients (37). Our analysis revealed that patients with tumors exhibiting high *KLC2* expression had a significantly lower probability of overall survival, indicating that elevated *KLC2* expression is associated with poor prognosis in CRC patients (n=1 061; p=0.031, **Figure 6F**). Taken together, these data demonstrate that the *miR-4745*-*KLC2* axis may serve as a prognostic indicator for overall survival in CRC.

## DISCUSSION

Colorectal cancer remains a major clinical challenge due to its high rates of chemoresistance and tumor recurrence, primarily driven by CSCs. Among potential therapeutic targets, miRNAs have gained significant attention for their pivotal role in regulating the CSC phenotype (38), offering new opportunities for targeting these cells in cancer therapy. In this study, we identified several miRNAs of interest that regulate the CSC phenotype, among which *miR-4745* was selected for further characterization. We demonstrated that *miR-4745* regulates CSC traits such as sphere-forming ability, self-renewal and chemoresistance. Mechanistically we identified the 3’ UTR of *KLC2* as a direct target of *miR-4745* and a key intermediate in the molecular mechanism controlling CSC traits.

Our bioinformatic analysis revealed that, in addition to its role in regulating *KLC2*, *miR-4745* indirectly downregulates the expression of genes that promote key hallmarks of cancer stem cells, including self-renewal (e.g. *SQSTM1* (39) and SPP1 (40)), invasion and metastasis (e.g. *MMP15* (41). This occurs in part through the activation of signaling pathways such as the Notch pathway (e.g. *HEY2,* (42)) or the ERK1/2 pathway (e.g. *GDF15*, (43), the modulation of the cellular stress response (e.g. *CYP2E1* (44)), and drug resistance (e.g., *LGMN* (45) and *UCP2* (46)). Importantly, we observed that *miR-4745* controls the expression of genes previously reported to negatively impact disease-free survival in colon cancer patients, including *GDF15*, *UCP2*, and *SPP1*. These data strongly suggest that, in addition to *KLC2*, *miR-4745* downregulates a network of critical downstream targets that may play a role in regulating the CSC phenotype.

*KLC2,* a light chain isoform of Kinesin-1, plays a critical role in the intracellular transport of various cargoes along microtubules. Mutations in the *KLC2* gene are associated with several neurological disorders, including spastic paraplegia, optic atrophy, and neuropathy (47). Notably, *KLC2* overexpression has been shown to promote radioresistance in non-small cell lung cancer (NSCLC), resulting in increased tumor volume and reduced survival in tumor-bearing mice. This suggests a potential role for *KLC2* in the survival of CSCs following radiotherapy. Additionally, elevated *KLC2* levels are correlated with decreased p53 phosphorylation and poor prognosis in NSCLC patients (48). *KLC2* is also a target of *miR-125b*, a microRNA involved in cancer cell migration and metastasis (49). Nonetheless, the target genes of *miR-125b* differ among various cancer types, in which its expression is frequently dysregulated. One known binding partner of *KLC2* is *SMAD2*, which is crucial for TGFβ signal transduction, leading to various effects during tumorigenesis. Further studies on the downstream pathways of *KLC2* could help uncover the molecular mechanisms behind its impact on CSC survival and its clinicopathological significance.

We observed a clear inverse correlation between *miR-4745* and *KLC2* mRNA levels across various stages of CRC cell lines, substantiating the existence of a *miR-4745/KLC2* axis. Furthermore, *KLC2* expression progressively increased with CRC advancement, peaking in stage III patient samples, suggesting it may serve as a potential local driver of CRC progression. However, further investigation is needed to understand the underlying reasons for the reduction in *KLC2* expression in stage IV tissues and metastatic sites.

Our study emphasizes the promise of targeting the *miR-4745/KLC2* pathway as a therapeutic strategy for colorectal cancer. To ensure that the observed effects on colorectal CSC traits were biologically relevant, and not simply the result of supraphysiological levels of *miR-4745-3p*, we quantified its expression by RT-qPCR in two CRC cell models transfected with decreasing concentrations of *miR-4745-3p* mimic. Although the effects were attenuated compared to transfection with 50 nM mimic, we still observed a measurable impact on the cells’ sphere-forming capacity at concentrations as low as 0.001 nM (**Supplementary Figure 4B**). Notably, this concentration aligns with the endogenous expression level of *miR-4745-3p* observed under control mimic conditions (**Supplementary Figure 4A**). These results suggest that the *miR-4745/KLC2* regulatory axis remains biologically relevant in CRC, even if its impact under physiological conditions may be moderate. Although *miR-4745-3p* exhibits low endogenous expression, it is likely that its regulation is context-dependent or limited to particular CRC subtypes or stem-like cell populations, analogous to other key miRNAs that play roles in stemness or stress response mechanisms.

Modulating the *miR-4745/KLC2* pathway may help reprogram CSC behavior, offering a strategy to overcome chemoresistance. For example, upregulating *miR-4745* or directly targeting *KLC2* could sensitize CSCs to chemotherapy, potentially reducing the likelihood of tumor recurrence and improving patient outcomes. While miRNA-based therapies faces challenges, particularly in achieving targeted and efficient systemic delivery, innovative technologies, including nanocapsules, nanocarriers, micro/nanoparticles, liposomes and PEGylated vesicles have been developed to address these limitations (50). Another promising alternative to modulate the *miR-4745/KLC2* axis could involve pharmacological agents that act as endogenous inducers or activators of *miR-4745*. In our previous study, we successfully induced *miR-148a-3p* expression with the anthelmintic drug niclosamide (16), demonstrating a practical strategy to enhance miRNA activity *in vivo* and raising the possibility of repurposing niclosamide as an adjuvant strategy to reduce CSC-mediated chemoresistance. Furthermore, assessing *miR-4745* expression in colorectal tumor samples, including those from chemotherapy-treated patients, could provide deeper insights into its role in therapy resistance. As mentioned, directly targeting *KLC2* through the development of a specific inhibitor or using PROTAC technology might represent another strategy to regulate the *miR-4745/KLC2* axis.

Finally, our miRnome-wide high content screening efforts identified other miRNAs, including *miR-497*, *miR-4723*, and *miR-4784*, which also contribute to reducing the CSC phenotype. Further investigation into the molecular mechanisms of these miRNAs could provide broader insights into CSC regulation and potential therapeutic strategies.

## CONCLUSIONS

In conclusion, our study identifies *miR-4745* as a novel and functionally significant regulator of colorectal CSC properties, acting in part through the direct repression of the kinesin light chain protein *KLC2*. We demonstrate that *miR-4745* impairs key CSC traits, including self-renewal, chemoresistance, and sphere-forming capacity, and delineate a broader regulatory network involving multiple downstream effectors implicated in CSC maintenance, tumor progression, and adverse clinical outcomes. The observed inverse correlation between *miR-4745* and *KLC2* expression across CRC stages reinforces the relevance of this regulatory axis in colorectal tumorigenesis. Although endogenously expressed at low levels, *miR-4745-3p* exhibits biologically meaningful activity within a physiologically relevant range, suggesting its potential as a context-specific modulator of CSC behavior. These findings provide a compelling rationale for therapeutic strategies aimed at restoring *miR-4745* expression or pharmacologically inhibiting *KLC2* to overcome CSC-driven chemoresistance. Finally, the identification of additional CSC-targeting miRNAs through our high-content screening broadens the scope for future combinatorial approaches. Altogether, our results highlight the therapeutic potential of miRNA-based interventions in targeting colorectal CSCs and improving patient outcomes.

## ABBREVIATIONS

*3′ UTR*: 3′ Untranslated region
5-FU: 5-fluorouracil
*ABCB1*: ATP-Binding Cassette Subfamily B Member 1
*ABCG2*: ATP-Binding Cassette G2
AcGFP: *Aequora coerulescens* Green fluorescent protein
*ALDH1A1*: Aldehyde Dehydrogenase 1A1
ATCC: American Type Culture Collection
*CCND3*: Cyclin D3
CRC: Colorectal cancer
CSCs: Cancer stem cells
*CYP2E1*: Cytochrome P450 2E1
dbDEMC: database of Differentially Expressed MiRNAs in human Cancers
DEAB: diethylaminobenzaldehyde
DEG: Differentially expressed genes
DMEM: Dulbecco’s Modified Eagle Medium
DS-Red: *Discosoma sp.* red fluorescent protein
EGF: Epidermal growth factor
ERK1/2: Extracellular Signal Regulated Kinase 1/2
FBS: Fetal bovine serum
FGF: Fibroblast growth factor
FSC: Forward Scatter
*GAPDH*: Glyceraldehyde-3-phosphate dehydrogenase
GDF15: Growth differentiation factor 15
GO: Gene Ontology
*HDAC4*: Histone Deacetylase 4
*HEY2*: Hes Related Family BHLH Transcription Factor With YRPW Motif 2
*KLC2*: Kinesin-light chain 2
*LGMN*: Legumain
mRNA: messenger RNA
miRNA: micro-RNA
*MIR31HG*: miR-31 Host Gene
*MMP15*: Matrix metalloproteinase 15
NSCLC: Non-Small Cell Lung Cancer
OE: overexpression
*PXR*: Pregnane X Receptor
QQ: quantile-to-quantile
RNA-seq: RNA sequencing
*RPLO*: Ribosomal protein L0
*RPL13*: Ribosomal protein L13
RPMI: Roswell Park Memorial Institute Medium
siRNA: small interfering RNA
*SMAD2*: SMAD Family Member 2
*SPP1*: Secreted Phosphoprotein 1
*SQSTM1*: Sequestosome 1
SSC: Side Scatter
TCGA: The Cancer Genome Atlas
TGF-β: Transforming growth factor-β
*UCP2*: Uncoupling Protein 2
WHO: World Health Organization

## AUTHORS’ CONTRIBUTIONS

C.P., A.D., J.M.P., and J.P. designed the research; C.P., A.D, K.L., J.M.P, and J.P. designed the experiments; P.B., K.L., J.R., O.B., G.P., A.B., S.M., C.M., E.P., L.B., C.D., A.A.G., and C.P. performed the experiments; P.B., K.L., J.R., O.B., G.P., A.B., S.M., C.M., E.P., L.B., C.D., A.A.G., E.R., J.P., J.M.P., A.D., and C.P. analyzed the data; C.P. K.L., and A.D. wrote the manuscript.

## ACKNOWLEDGMENTS

We thank C. Duperray and F. Leccia (IRMB, Montpellier) from the Montpellier RIO Imaging platform for flow cytometry experiments.

## DECLARATION

We confirm that all research involving human subjects, human material, or human data in our study was conducted in full accordance with the ethical principles outlined in the Declaration of Helsinki.

### Cell lines and patient-derived tumor cell cultures

Patient-derived colon cancer cell lines (CPP1, CPP6, CPP19, CPP25, CPP36, and CPP44) and circulating tumor cell lines (CTC44 and CTC45) were established from colorectal cancer biopsies and peripheral blood samples collected at CHU-Carémeau (Nîmes, France) as part of a previously clinical study (20,21), registered on ClinicalTrials.gov (Identifier NCT01577511 ; Principal Investigator: Dr. Jean-François Bourgaux ; First submitted: 2012-04-12; First submitted that met QC criteria : 2012-04-13 ; First posted (estimated) : 2012-04-16 ; Last Update submitted that met QC criteria : 2021-06-18 ; Last update posted: 2021-06-21; Last verified : 2020-01 ; https://clinicaltrials.gov/study/NCT01577511).

The study protocol was approved by the French Ethics Committee “Comité de Protection des Personnes Sud Méditerranée III (authorization number 2011-A01141-40),” which operates in full compliance with the Declaration of Helsinki. Written informed consent was obtained from all participants involved in the study, in accordance with all relevant ethical regulations. Commercial CRC cell lines used in this study - SW480 (RRID: CVCL_0546), SW620 (RRID:CVCL_0547), HT29 (RRID:CVCL_A8EZ), LS174T (RRID:CVCL_1384), HCT116 (RRID:CVCL_0291), LoVo (RRID:CVCL_0399), and Colo205 (RRID:CVCL_0218) were obtained from ATCC and used in accordance with standard ethical practices and institutional guidelines.

### Colon tumor tissue samples

RNA extracted from 41 colon tumor tissue samples was obtained through the Clinical and Biological Database BCBCOLON (Institut du Cancer de Montpellier—Val d’Aurelle, France) registered at ClinicalTrials.gov (Identifier #NCT03976960 ; Principal Investigator: Dr. Eric Assenat ; First submitted: 2019-06-05; First submitted that met QC criteria : 2019-06-05 ; First posted : 2019-06-06 ; Last Update submitted that met QC criteria : 2025-02-11 ; Last update posted: 2025-02-12; Last verified : 2025-02 ; https://clinicaltrials.gov/study/NCT03976960).

These samples were collected in compliance with French legislation and declared to the French Ministry of Higher Education and Research (Declaration No. DC-2008–695). The study received approval from the French Ethics Committee (CPP Sud Méditerrannée III, Ref#2014.02.04) and the ICM Translational Research Committee (ICM-CORT-2018-28). All participants provided written consent, and all relevant ethical regulations were strictly followed.

---

**Supplementary Figure 2:**
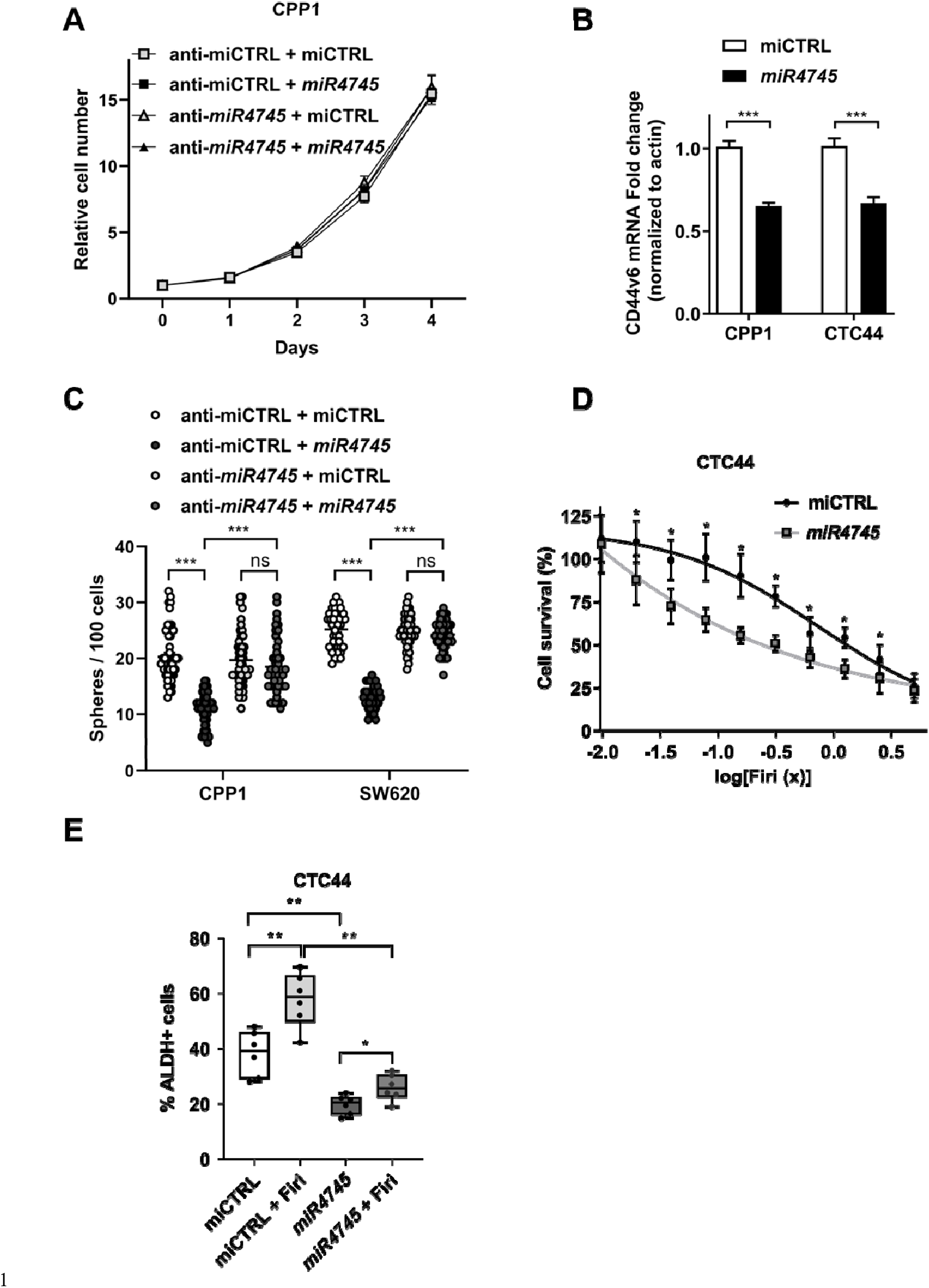
Overexpression of *miR-4745* represses the CSC phenotype in vitro. **(a)** Cell proliferation was assessed daily for 4 days using the sulforhodamine B colorimetric assay in patient-derived CPP1 cells co-transfected for 72 hours with either anti-*miR-4745* or anti-miCTRL, along with *miR-4745* or miCTRL mimics. **(b)** CD44v6 mRNA expression in patient-derived CPP1 colon cancer cells and the circulating tumor cell line (CTC44), transfected with *miR-4745* or a negative control mimic (miCTRL). Data are presented as mean ± SEM (n=3). Results are shown as fold change relative to cells transfected with the negative control mimic. **(c)** Percentage of sphere-forming cells in patient-derived CPP1 and SW620 colon cancer cells co-transfected for 72 hours with either anti-*miR-4745* or anti-miCTRL, along with *miR-4745* or miCTRL mimics. Individual replicate values (n=15 per experiment) are plotted along with the mean ± SEM of three independent experiments. **(d)** Percentage of surviving CSCs (Aldefluor-positive cells) 48 hours after exposure to the indicated concentrations of FIRI (1X = 50µM 5-FU + 500nM SN38) in vitro. Sorted CTC44 Aldefluor-positive cells were transfected with 50nM *miR-4745-3p* (miR-4745) or miRVana microRNA Mimic Negative Control #1 (miCTRL) mimics, 24hours before FIRI treatment. Data are presented as mean ± SEM (n = 3). **(e)** Percentage of Aldefluor-positive cells (% ALDH^+^ cells) in CTC44 cells transfected with miR-4745 or miCTRL mimics and then exposed to chemotherapy (FIRI = 5µM 5-FU + 50nM SN38) for 72 hours. Data are presented as mean ± SEM (n = 6). *, p < 0.05; **, p < 0.005; ***, p < 0.001; ns = not significant, Mann-Whitney test.

**Supplementary Figure 4:**
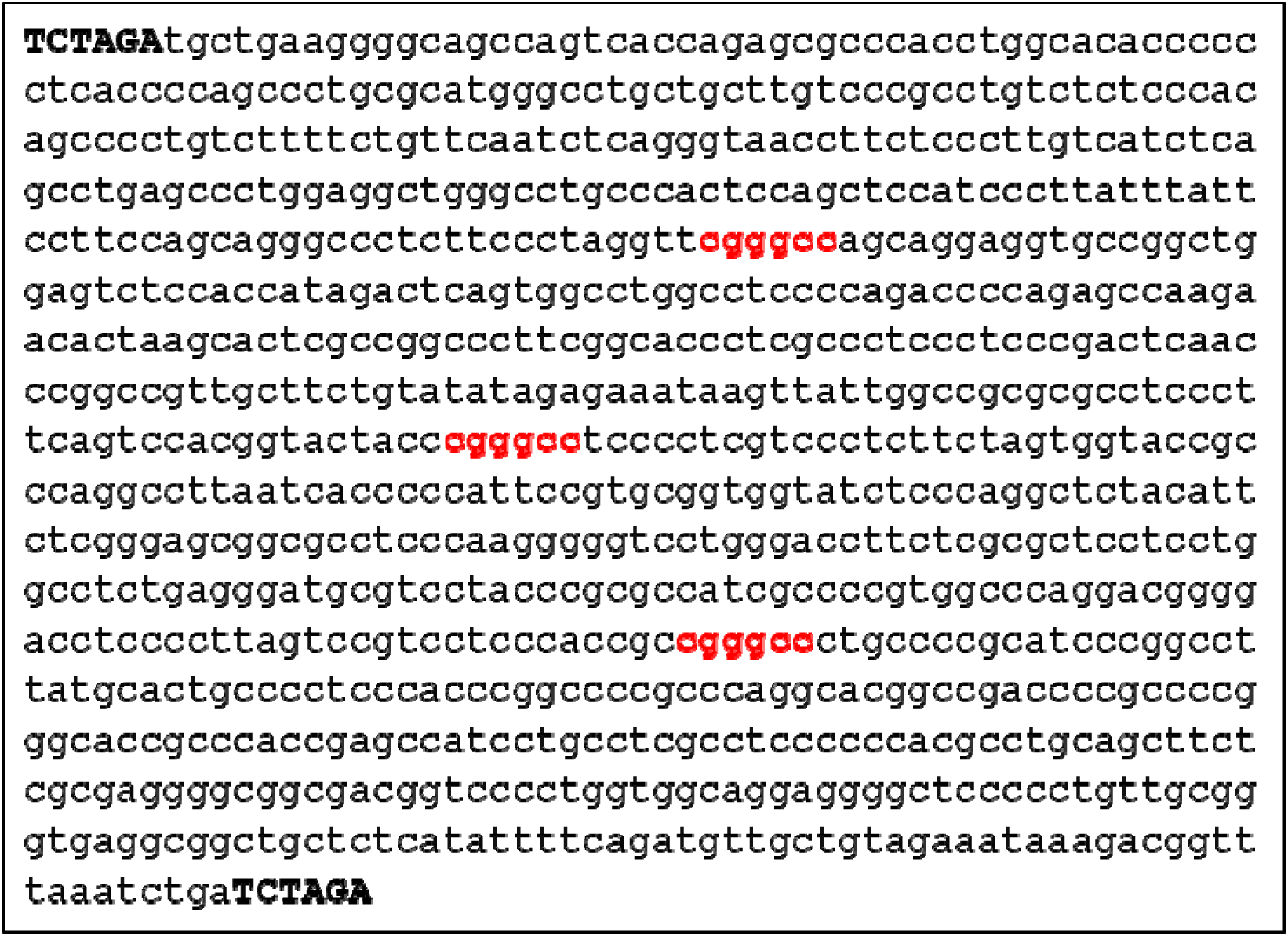
*KLC2* 3’UTR sequence. *KLC2* 3’UTR sequence with XbaI sites (highlighted at the ends) cloned into the pmiR-sensor plasmid. The three predicted binding sites for *miR-4745* are highlighted in red.

**Supplementary Figure 5:**
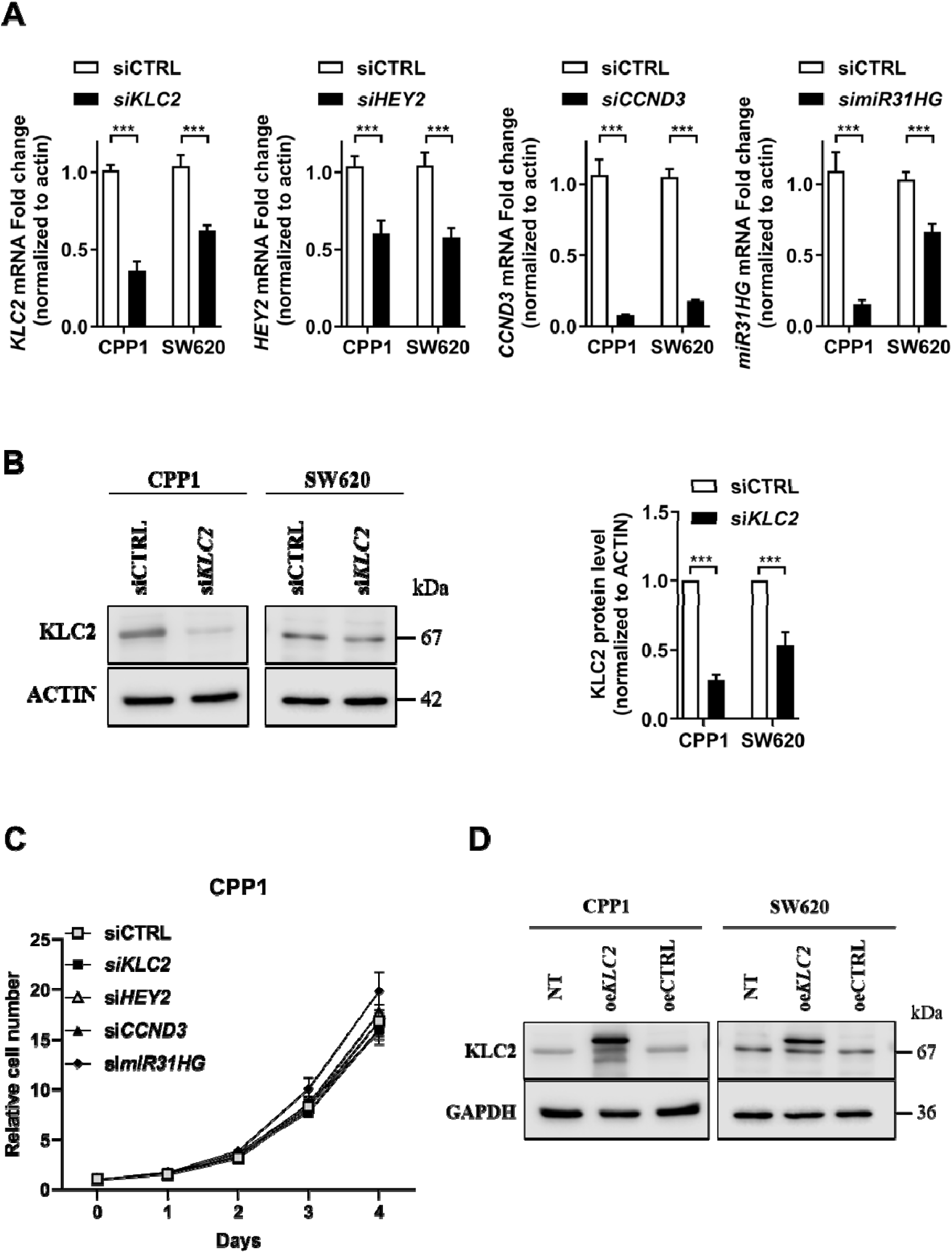
*KLC2* inactivation inhibits colorectal CSC phenotype. **(a)** mRNA expression levels of *KLC2*, *HEY2*, *CCND3*, and *miR31HG* in CPP1 and SW620 cells transfected for 72 hours with 50nM control siRNA (siCTRL) or specific siRNA targeting *KLC2* (siKLC2), *HEY2* (siHEY2), *CCND3* (siCCND3), or *miR31HG* (simiR31HG). Data are presented as mean ± SEM (n=3) and are expressed as fold change relative to siCTRL-transfected cells. **(b)** Representative Western blot analysis (left panel) and quantification (right panel) of KLC2 protein levels in patient-derived CPP1 and SW620 colon cancer cells transfected for 72 hours with 50nM control siRNA (siCTRL) or siRNA targeting KLC2 (siKLC2). Data are presented as mean +- SEM (n=4). **(c)** Cell proliferation was assessed daily for 4 days using the sulforhodamine B colorimetric assay in patient-derived CPP1 cells transfected for 72 hours with 50nM control siRNA (siCTRL) or specific siRNAs targeting *KLC2* (siKLC2), *HEY2* (siHEY2), *CCND3* (siCCND3), or *miR31HG* (simiR31HG). Data are presented as mean +/- SEM (n=3). ***p < 0.001; Mann-Whitney test. **(d)** Western blot analysis of KLC2 protein levels (both endogenous KLC2 and KLC2-3x Flag) in patient-derived CPP1 and SW620 colon cancer cells transfected with a KLC2-overexpressing plasmid (oe-KLC2) or an empty vector (oe-CTRL). NT refers to non-transfected cells.

**Supplementary Figure 6:**
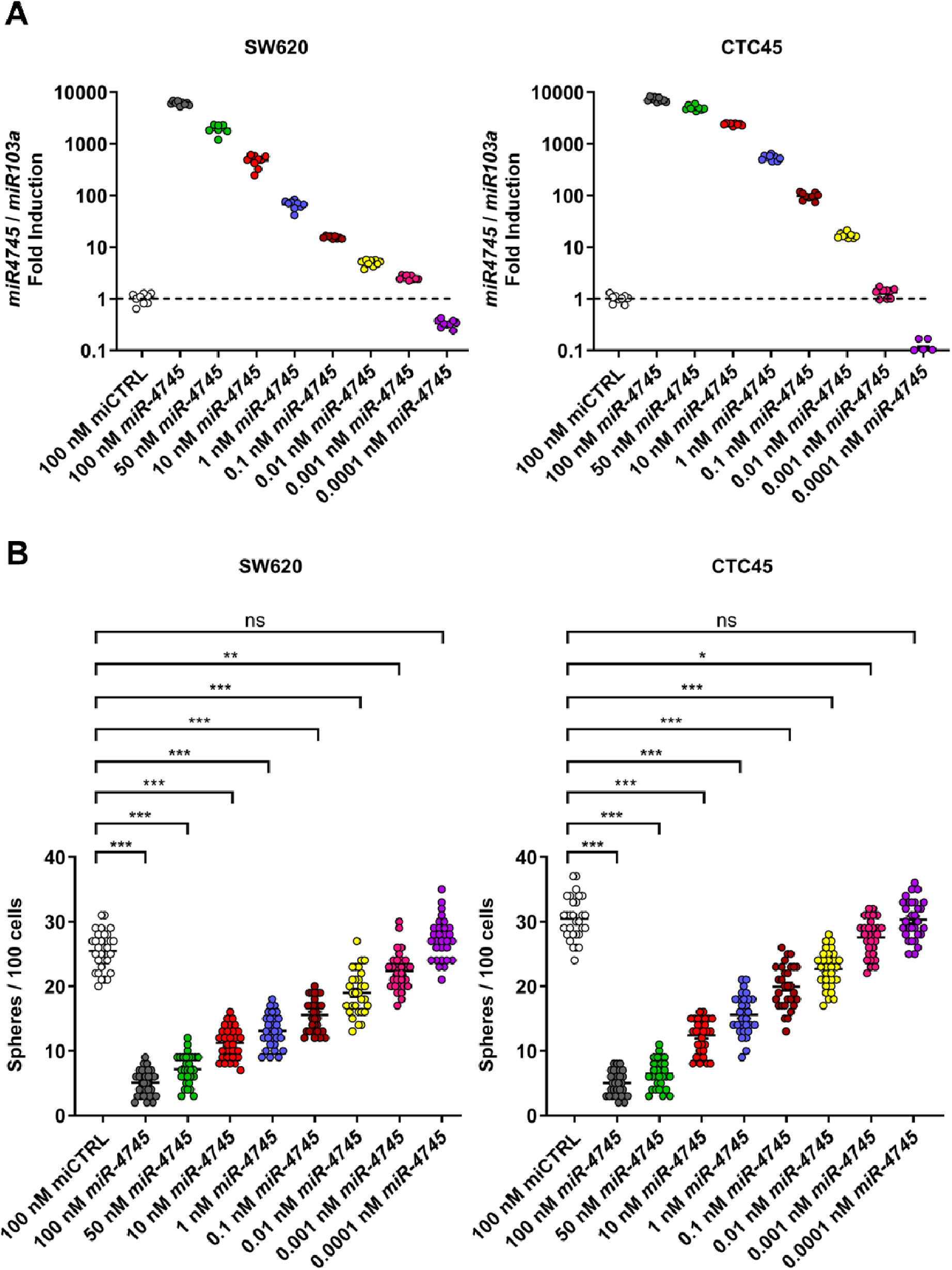
*miR-4745-3p* mimic induces dose-dependent changes in expression levels and sphere-forming capacity in CRC cell lines. **(a)** Relative expression levels of *miR-4745-3p* in SW620 colon cancer cells and in the circulating tumor cell line CTC45, transfected for 72 hours with varying concentrations of *miR-4745-3p* or control (miCTRL) mimics. Results are expressed as fold change relative to miCTRL-transfected cells and presented as mean ± SEM (n = 3). **(b)** Sphere-forming efficiency of SW620 and CTC45 cells transfected for 72 hours with varying concentrations of miR-4745-3p or miCTRL mimics. Data represent the percentage of sphere-forming cells (mean ± SEM; n = 3 independent experiments, 10 replicates per experiment). Statistical significance was assessed using one-way ANOVA: *, p < 0.05, **, p < 0.005, ***, p < 0.001, ns = not significant.

**Supplementary Table S1:**
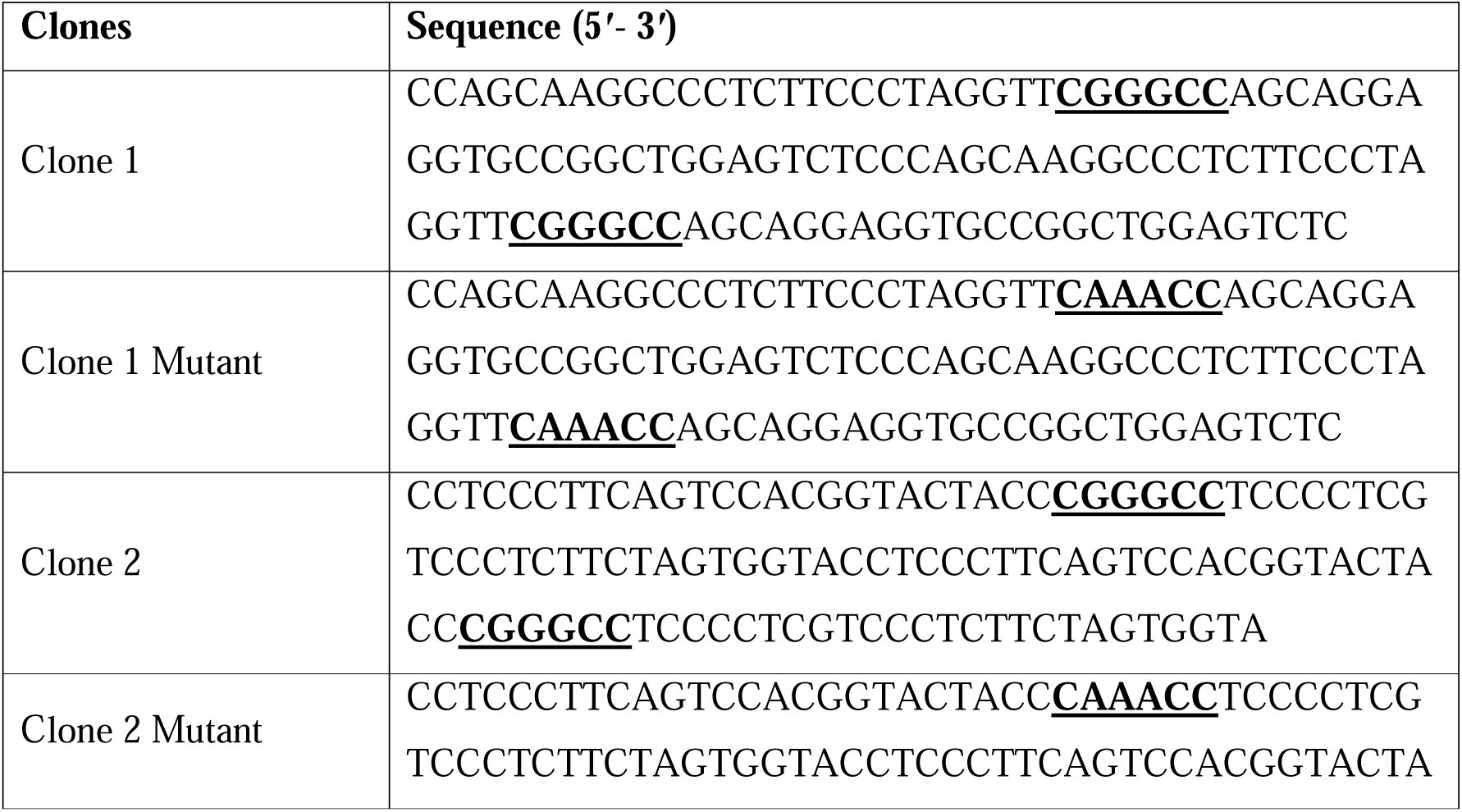
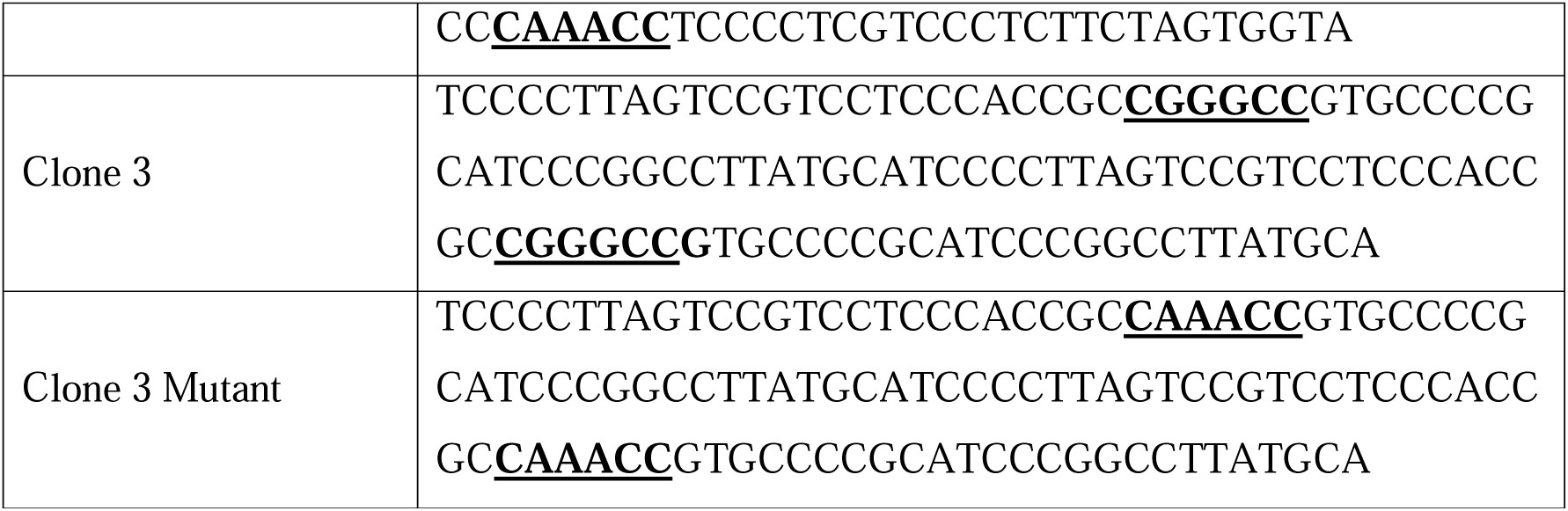
*KLC2* 3’-UTR regions cloned into the pmiR-sensor plasmid. The underlined region indicates the predicted binding sequence for the seed region of *miR-4745*.

**Supplementary Table S2:**
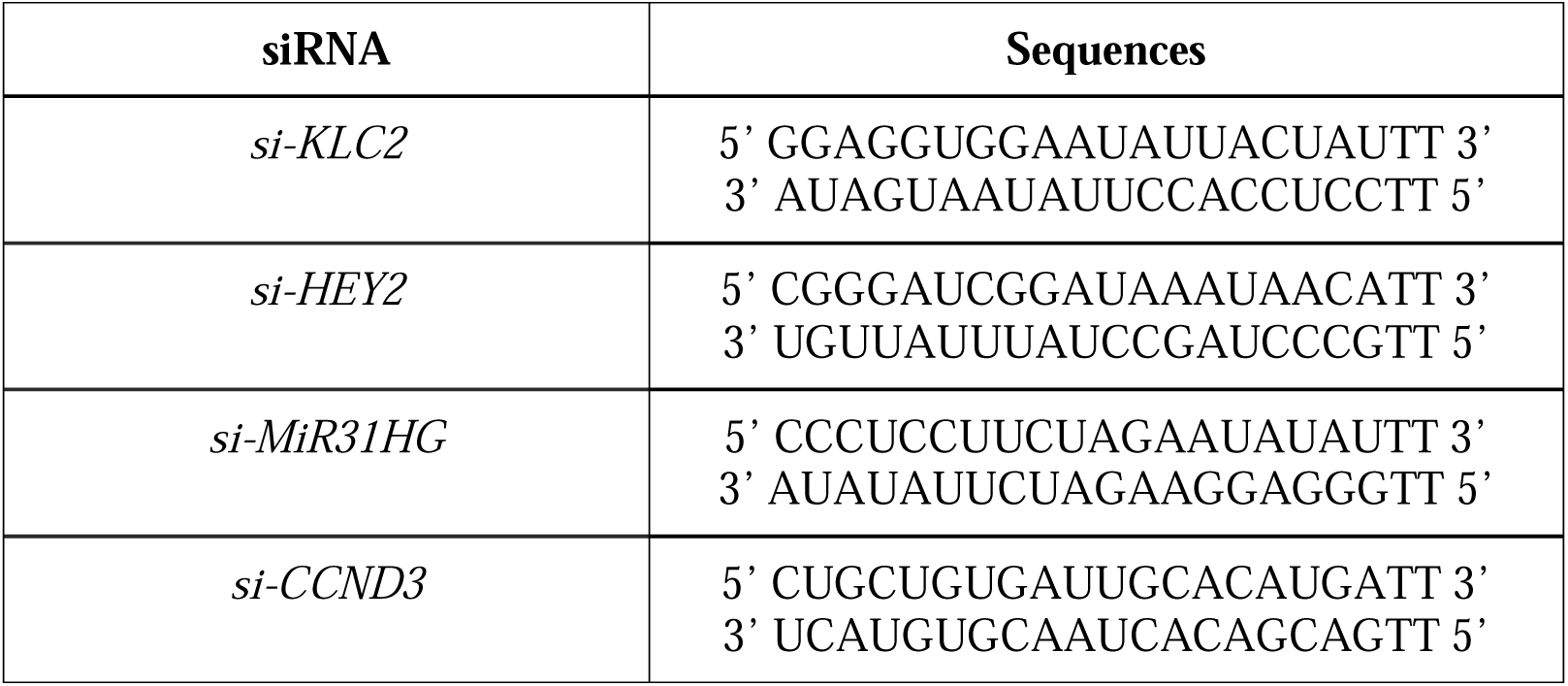
List of siRNA duplexes targeting *KLC2*, *HEY2*, *CCND3*, and *miR31HG*.

**Supplementary Table S3:**
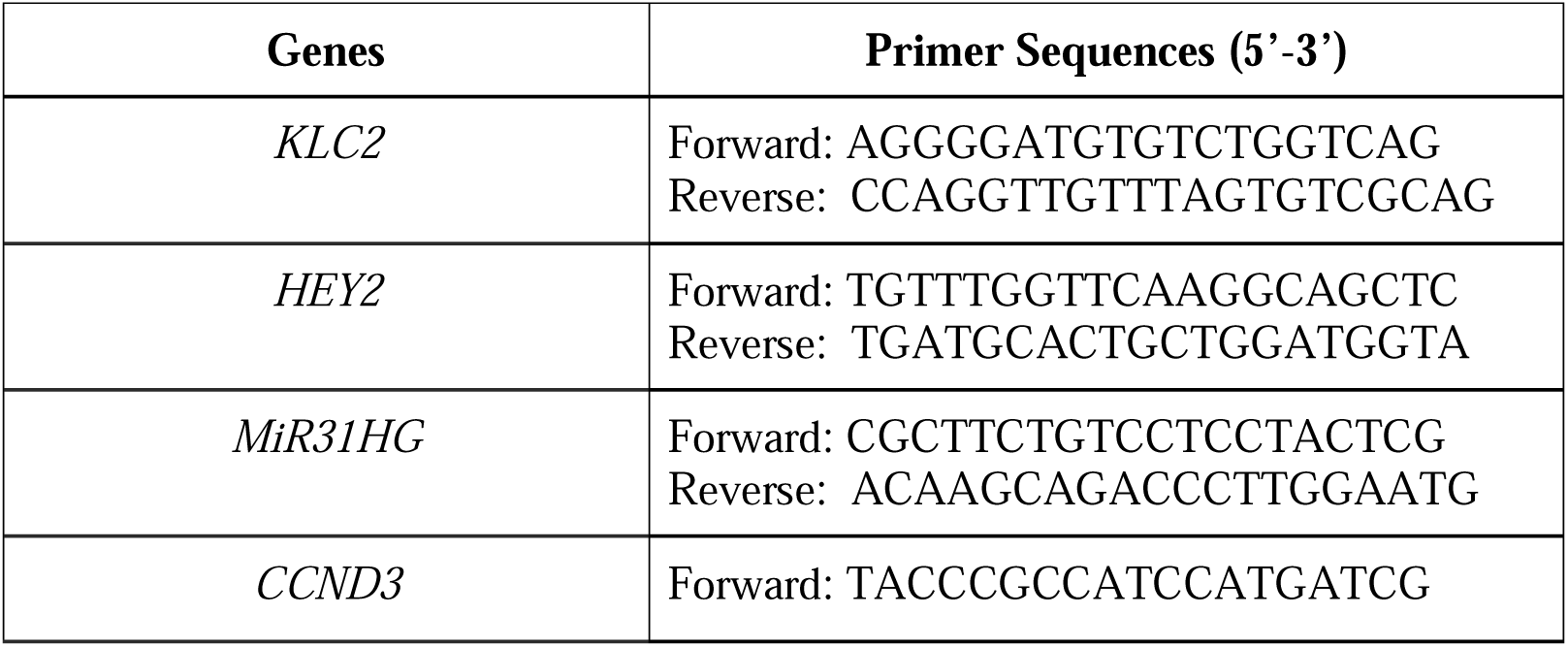
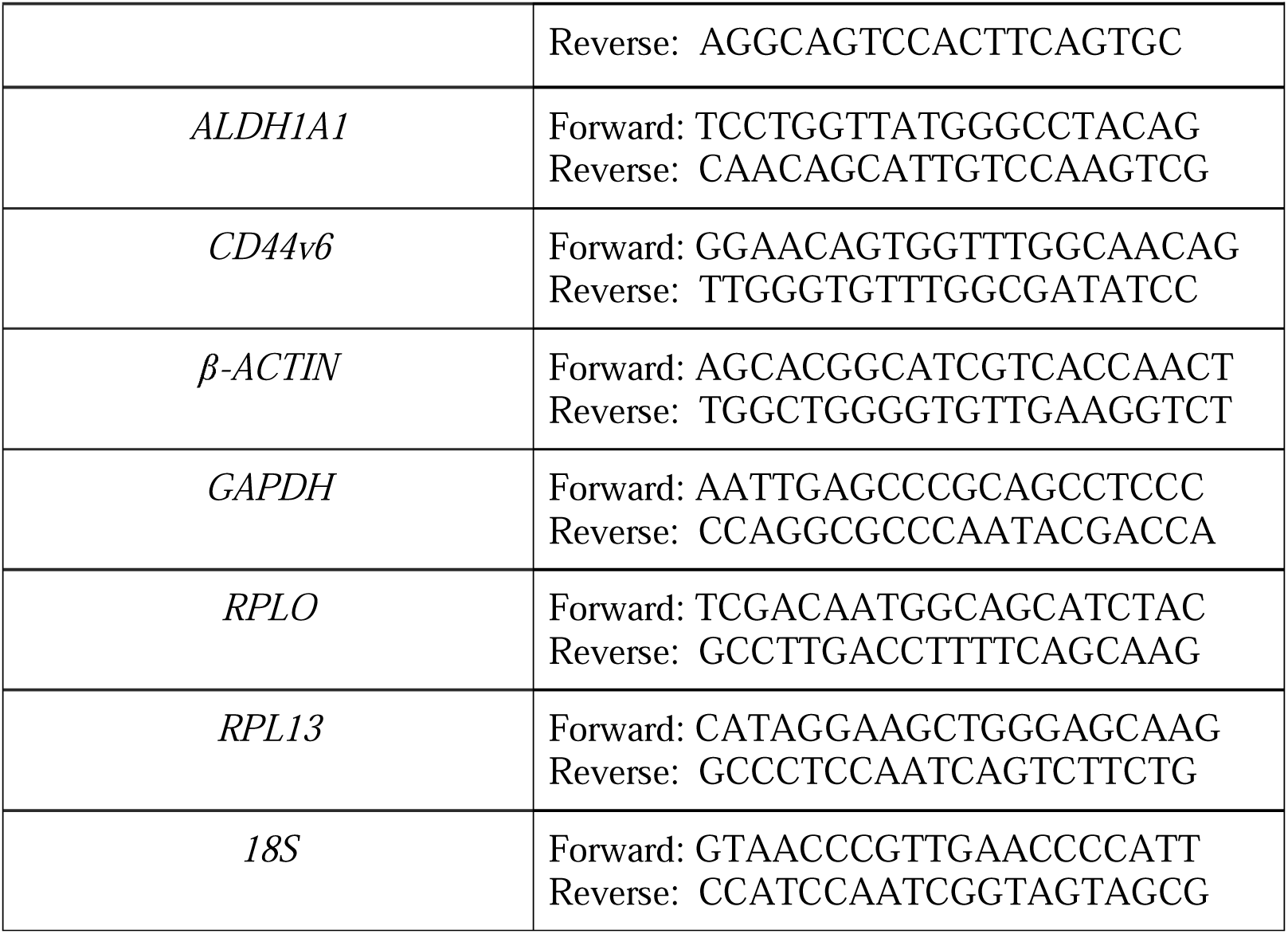
Primer sequences.

**<u>Supplementary Table S4:</u> List of miRNAs identified from human miRNome-wide screening using a miRNA overexpression (OE) library that deregulate ALDEFLUOR activity in the SW620 cell line model (Excel file).**

**<u>Supplementary Table S5:</u> List of transcripts that are downregulated or upregulated in the transcriptome analysis following *miR-4745* overexpression in CPP1 cells.**

**<u>Supplementary Table S6:</u> List of transcripts that are downregulated or upregulated in the translatome analysis, specifically in the highly translated fractions (heavy fractions with more than 3 ribosomes per mRNA).**

